# Mouse models of *SYNGAP1*-related intellectual disability

**DOI:** 10.1101/2023.05.25.542312

**Authors:** Yoichi Araki, Elizabeth E. Gerber, Kacey E. Rajkovich, Ingie Hong, Richard C. Johnson, Hey-Kyoung Lee, Alfredo Kirkwood, Richard L. Huganir

**Author notes:** Corresponding author Richard Huganir. These authors contributed equally (Yoichi Araki, Elizabeth E. Gerber, Kacey E. Rajkovich, Ingie Hong).

## Abstract

SYNGAP1 is a Ras-GTPase activating protein highly enriched at excitatory synapses in the brain. *De novo* loss-of-function mutations in *SYNGAP1* are a major cause of genetically defined neurodevelopmental disorders (NDD). These mutations are highly penetrant and cause *SYNGAP1*-related intellectual disability (SRID), a NDD characterized by cognitive impairment, social deficits, early-onset seizures, and sleep disturbances (1-5). Studies in rodent neurons have shown that Syngap1 regulates developing excitatory synapse structure and function (6-11), and heterozygous *Syngap1* knockout mice have deficits in synaptic plasticity, learning and memory, and have seizures (9, 12-14). However, how specific *SYNGAP1* mutations found in humans lead to disease has not been investigated in vivo. To explore this, we utilized the CRISPR-Cas9 system to generate knock-in mouse models with two distinct known causal variants of SRID: one with a frameshift mutation leading to a premature stop codon, *SYNGAP1; L813RfsX22,* and a second with a single-nucleotide mutation in an intron that creates a cryptic splice acceptor site leading to premature stop codon, *SYNGAP1; c.3583-9G>A*. While reduction in *Syngap1* mRNA varies from 30-50% depending on the specific mutation, both models show ∼50% reduction in Syngap1 protein, have deficits in synaptic plasticity, and recapitulate key features of SRID including hyperactivity and impaired working memory. These data suggest that half the amount of SYNGAP1 protein is key to the pathogenesis of SRID. These results provide a resource to study SRID and establish a framework for the development of therapeutic strategies for this disorder.

**Significance Statement:** SYNGAP1 is a protein enriched at excitatory synapses in the brain that is an important regulator of synapse structure and function. *SYNGAP1* mutations cause *SYNGAP1*-related intellectual disability (SRID), a neurodevelopmental disorder with cognitive impairment, social deficits, seizures, and sleep disturbances. To explore how *SYNGAP1* mutations found in humans lead to disease, we generated the first knock-in mouse models with causal SRID variants: one with a frameshift mutation and a second with an intronic mutation that creates a cryptic splice acceptor site. Both models show decreased *Syngap1* mRNA and Syngap1 protein and recapitulate key features of SRID including hyperactivity and impaired working memory. These results provide a resource to study SRID and establish a framework for the development of therapeutic strategies.

**Highlights:**

1. Two mouse models with *SYNGAP1*-related intellectual disability (SRID) mutations found in humans were generated: one with a frameshift mutation that results in a premature stop codon and the other with an intronic mutation resulting in a cryptic splice acceptor site and premature stop codon.
2. Both SRID mouse models show 35∼50% reduction in mRNA and ∼50% reduction in Syngap1 protein.
3. Both SRID mouse models display deficits in synaptic plasticity and behavioral phenotypes found in people.
4. RNA-seq confirmed cryptic splice acceptor activity in one SRID mouse model and revealed broad transcriptional changes also identified in *Syngap1^+/-^* mice.
5. Novel SRID mouse models generated here provide a resource and establish a framework for development of future therapeutic intervention.

## Introduction

Recent advances in next-generation sequencing technology have led to the discovery of numerous causative genetic mutations in neurodevelopmental disorders (NDDs) (1, 2, 4, 5). Many of these mutations are found in genes that encode proteins at glutamatergic synapses, or those that regulate them, including post-synaptic density (PSD) components such as SH3 and multiple ankyrin repeat domains 3 (SHANK3), Postsynaptic density protein-95 (PSD-95), α- amino-3-hydroxy-5-methyl-4-isoxazolepropionic acid (AMPA) receptors, Neuroligins, presynaptic components and channels including Syntaxin-binding protein 1 (STXBP1), Sodium channel protein type 1 subunit alpha 1 (SCN1A) and 2 (SCN2A), Neurexins, and transcription factors such as AT-rich interactive domain-containing protein 1B (ARID1B), Chromodomain-helicase-DNA-binding protein 2 (CHD2), and 8 (CHD8) (2). These proteins serve essential regulatory and/or structural functions required for synaptic transmission and plasticity at glutamatergic synapses (3), strongly implicating these processes in disease pathogenesis of many NDDs. Among these, *SYNGAP1* is one of the most frequently mutated genes in NDDs (2, 4, 5). In one large exome sequencing study in the UK, *SYNGAP1* was the fourth-most prevalent mutated gene in NDD and found in 0.7% of the NDD population (4).

*SYNGAP1* encodes SYNGAP1, a Ras-GTPase activating protein (GAP) that is one of the most abundant components of the post-synaptic region in glutamatergic neurons. SYNGAP1 facilitates GTP hydrolysis to GDP through its GAP domain and thereby negatively regulates Ras activity (15, 16). SYNGAP1 is required for a major form of synaptic plasticity, long-term potentiation (LTP), a process thought to be critical to learning and memory (6-9). During LTP, SYNGAP1 is rapidly phosphorylated by CaMKII and is dispersed from the synapses by NMDAR activity (10, 11). This rapid dispersion triggers synaptic Ras activity, AMPAR insertion, and synaptic spine enlargements (11). SYNGAP1 has also been proposed to play a structural role in the post-synaptic density as it undergoes liquid-liquid phase separation (LLPS) through its interaction with PSD-95 (17).

Multiple start sites and alternative splice sites in *Syngap1* allow for several protein isoforms (***SI Appendix*, Fig. S1A**) that differ in structure, function, and temporospatial expression (18-23). N-terminal isoforms include A1/2/3/4, B, C, D, and E (21-23). In mouse cortex, mRNA levels of A and B present early in development at postnatal day 4 are similar to those in adult mice, while mRNA levels of C are much lower. Interestingly, A and B reach two to three times adult levels in mice at postnatal day 14 before diminishing again (23). There are four C-terminal isoforms that have been identified: α1, α2, β, and γ. The β isoform is the dominant isoform early in postnatal development, whereas α2 is the most abundant isoform at later stages. α1 is expressed at low levels during early postnatal development and increases to become the second most abundant isoform by postnatal day 42. Subcellularly, α1 is highly enriched at excitatory synapses, while the β isoform is more often in localized to the cytosol (20).

Mutations in *SYNGAP1* cause *SYNGAP1*-related intellectual disability (SRID). SRID is characterized by neurodevelopmental delay and mild-to-severe intellectual disability (24-28). About 80 to 85% of individuals with SRID have comorbid epilepsy, a subset of which have myoclonic astatic epilepsy (Doose syndrome) or epilepsy with myoclonic absences (29). Some people with SRID also have strabismus and hypotonia with significant motor deficits (28). The occurrence of autism spectrum disorder (ASD) is estimated to be as high as 50% in SRID, and includes stereotypic behaviors, obsessions with certain objects, and social deficits. Poor attention, impulsivity, lack of self-preservation instinct, self-directed and other-directed aggressive behavior, elevated pain threshold, hyperacusis, and sleep disorders have also been observed (30-33). Currently, treatment for SRID is limited to physical and behavioral therapy and specialty consultations for various symptoms. No standardized guidelines are available regarding the choice of specific anti-seizure medications. While some individuals with SRID are medication-responsive, at least half are treatment-resistant (29).

Heterozygous *Syngap1* knockout mice recapitulate several phenotypes of SRID including deficits in learning, memory, social behavior, as well as hyperactivity, repetitive behavior, and seizures (9, 12-14). However, to date, there have been no in vivo studies of SRID pathogenesis with mouse models harboring known causal variants found in humans. To investigate the pathophysiology of SRID with the aim to open new avenues of therapeutic investigation, we generated two knock-in mouse models with known causal variants of SRID including a frameshift mutation which leads to a premature stop codon, *SYNGAP1; L813RfsX22* (24) and a single-nucleotide mutation in an intron that creates a cryptic splice acceptor site and a premature stop codon, *SYNGAP1; c.3583-9G>A* (34).

## Results

### Generation of knock-in *Syngap1* mutant mice

To investigate the consequences of SRID *SYNGAP1* mutations in vivo, we used the CRISPR-Cas9 system to generate two knock-in mouse models. For one model, we selected a *de novo* frameshift mutation that leads to a premature stop codon (*SYNGAP1*-L813RfsX22). Importantly, the individual carrying this mutation has been well-characterized clinically and has classic features of SRID including developmental delay, intellectual disability, and epilepsy (24) (**Fig. 1*A***; Left panel). For our second knock-in mouse line, we selected a *de novo SYNGAP1* mutation also associated with SRID that is within an intron and creates a cryptic splice site and a premature stop codon (c.3583-9 G>A) (34) (**Fig. 1*A***; Right panel). To introduce mutations, Guide RNA, Cas9, and homology-directed repair DNA (HDR-DNA) oligos were injected into fertilized eggs (**Fig. 1*B***). Additional restriction enzyme sites were introduced for screening purposes. Newborn mice were screened by PCR and restriction enzyme digest. The introduction of mutations was confirmed by Southern blot (**Fig. 1*C***), Sanger sequencing of genomic DNA (**Fig. 1*D***), and Sanger sequencing of cDNA from heterozygous mice (**Fig. 1*E***). We confirmed that the cryptic splicing mutation (c.3583-9G>A) caused aberrant splicing resulting in a 7-base pair(bp) addition within mRNA transcripts (**Fig. 1*E:*** Right panel, red box). In both L813RfsX22 and c.3583-9G>A lines, the chromatogram peaks of the mutant allele were consistently lower, indicating that Syngap1 mutant mRNA was unstable or inefficiently transcribed.

**Figure 1.**
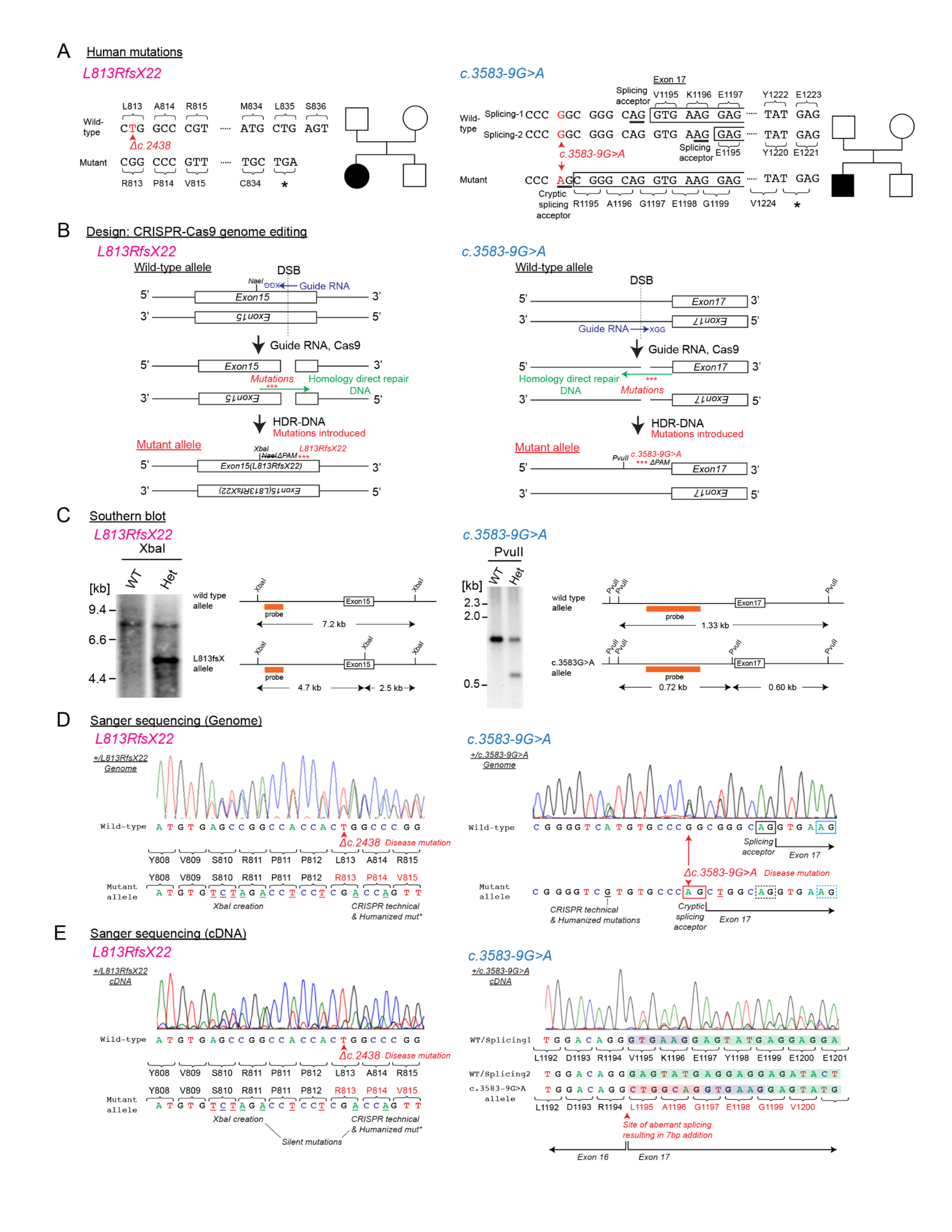
**Generation of *SYNGAP1*-related intellectual disability (SRID) model mice** (A) Schematic of SRID mutations. *SYNGAP1; L813RfsX22*, a frameshift mutation leading to a premature stop codon. *SYNGAP1; c.3583-9G>A*, a single nucleotide mutation in an intron that creates a cryptic splice acceptor site, the addition of 17 bp at the end of exon 17, and a premature stop codon. Pedigrees of affected individuals are shown to the right of each mutation. (B) CRISPR/Cas9 gene-engineering for SRID model mice. Guide RNA, Cas9 protein, and homology-directed repair DNA were injected into fertilized egg. (C) Southern blot confirming mutations in SRID model mice. The correct overall structure of the genome was confirmed by Southern blotting. The autoradiogram images of Southern blotting both from wild-type (WT) and heterozygotes (+/L813RfsX22 or c.3583-9G>A) are shown. Mutant allele carrying XbaI site (L813RfsX22) and PvuII site (c.3583-9G>A) was cut by restriction enzyme to confirm the appearance of correct size DNA. (D) Genomic DNA isolated from heterozygous SRID mice was sequenced by Sanger sequencing. The electropherogram near the disease mutation site is shown. SRID mutations, restriction enzyme sites for screening, and protospacer adjacent motif (PAM) site deletions are shown. Underline denotes the CRISPR technical mutations. Rectangular boxes indicate the splicing acceptor site (Black: splicing-1, Cyan: splicing-2, Red: cryptic splicing site emerged by mutation, Dotted: obsolete by disease mutation). (E) cDNA isolated from heterozygous SRID mice was sequenced by Sanger sequencing. The electropherogram near the disease mutation site is shown. In *Syngap1^+/L813Rfsx22^* mice, sequencing confirmed SRID mutation L813RfsX22, XbaI site creation for screening, and PAM site deletions. Sequencing of cDNA from *Syngap1^+/c.3583-9G>A^* mice confirmed the aberrant 7bp addition at the beginning of Exon 17 (Red boxes). Blue boxes indicate the 6bp difference between splicing-1 and splicing-2. Underline denotes the CRISPR technical mutations.

### *Syngap1^+/L813RfsX22^* mice show 30-40 % less *Syngap1* mRNA and 50% less SYNGAP1 protein

Northern blotting of whole brain mRNA extracts from *Syngap1^+/L813Rfsx22^* mice revealed a 30-40% reduction of mRNA (62.3 ± 4.1 % (mean ± S.E.M.) in comparison to wild-type controls, suggesting that mutant *Syngap1* mRNA is subject to nonsense-mediated decay (NMD) (35) as expected due to the premature stop codon (**Fig. 2*A***). This result was confirmed by quantitative PCR (qPCR) (69.9% ± 8.0 % compared to wild-type) (**Fig. 2*B***). Western blotting showed Syngap1 protein was significantly diminished in whole brain from *Syngap1^+/L813RfsX22^* mice (50.7 ± 3.5 % compared to wild-type, p < 0.0001 ***), similar to that in heterozygous *Syngap1* knock-out mice, *Syngap1^+/-^* (52.0 ± 9.7 % compared to wild-type, p = 0.0004 ***) (**Fig. 3*A***).

**Figure 2.**
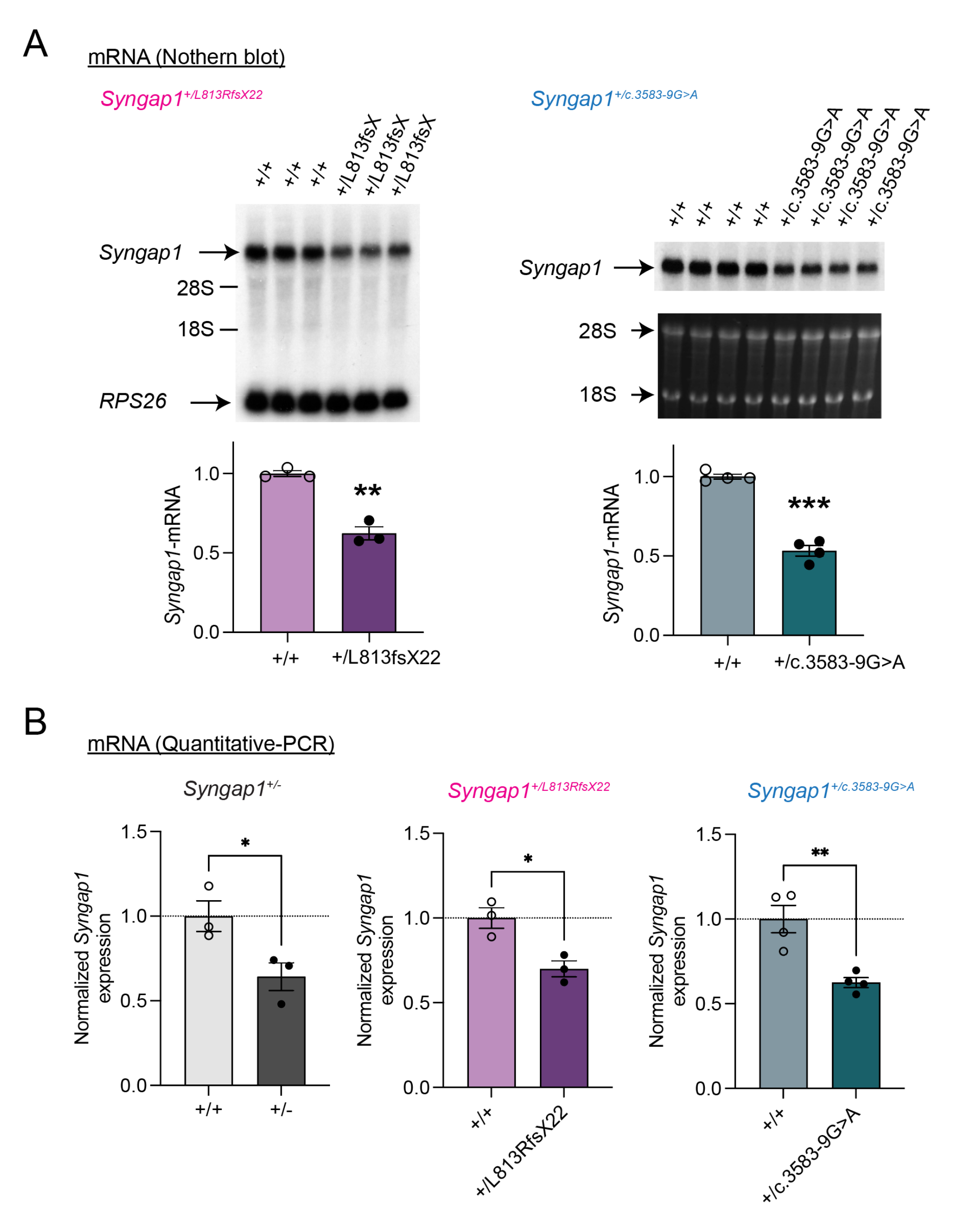
**Expression of mRNA from SRID disease model mice was significantly reduced by 30-50%** (A) Northern blotting of mRNAs from SRID mice. The autoradiogram of northern blotting using total brain of wild-type (+/+) or heterozygous mice (+/L813RfsX22 or +/c.3583-9G>A) was shown. L813RfsX22; *Rps26* bands were used for loading controls. The amount of *Syngap1* mRNA was normalized by *Rps26* mRNA levels. C.3583-9G>A; 28S and 18S ribosomal RNA was used for loading controls. SYNGAP1 mRNA was normalized by ribosomal RNA levels. Two-tailed t-test was performed (* p< 0.05, ** p< 0.01, *** p<0.001). (B) Quantitative-PCR from SRID mice and *Syngap1*^+/-^ mice. The mRNA quantification normalized to actin expression is shown. All mutant mice show 30-40% *Syngap1* mRNA reduction. Two-tailed t-test was performed (* p< 0.05, ** p< 0.01, *** p<0.001).

**Figure 3.**
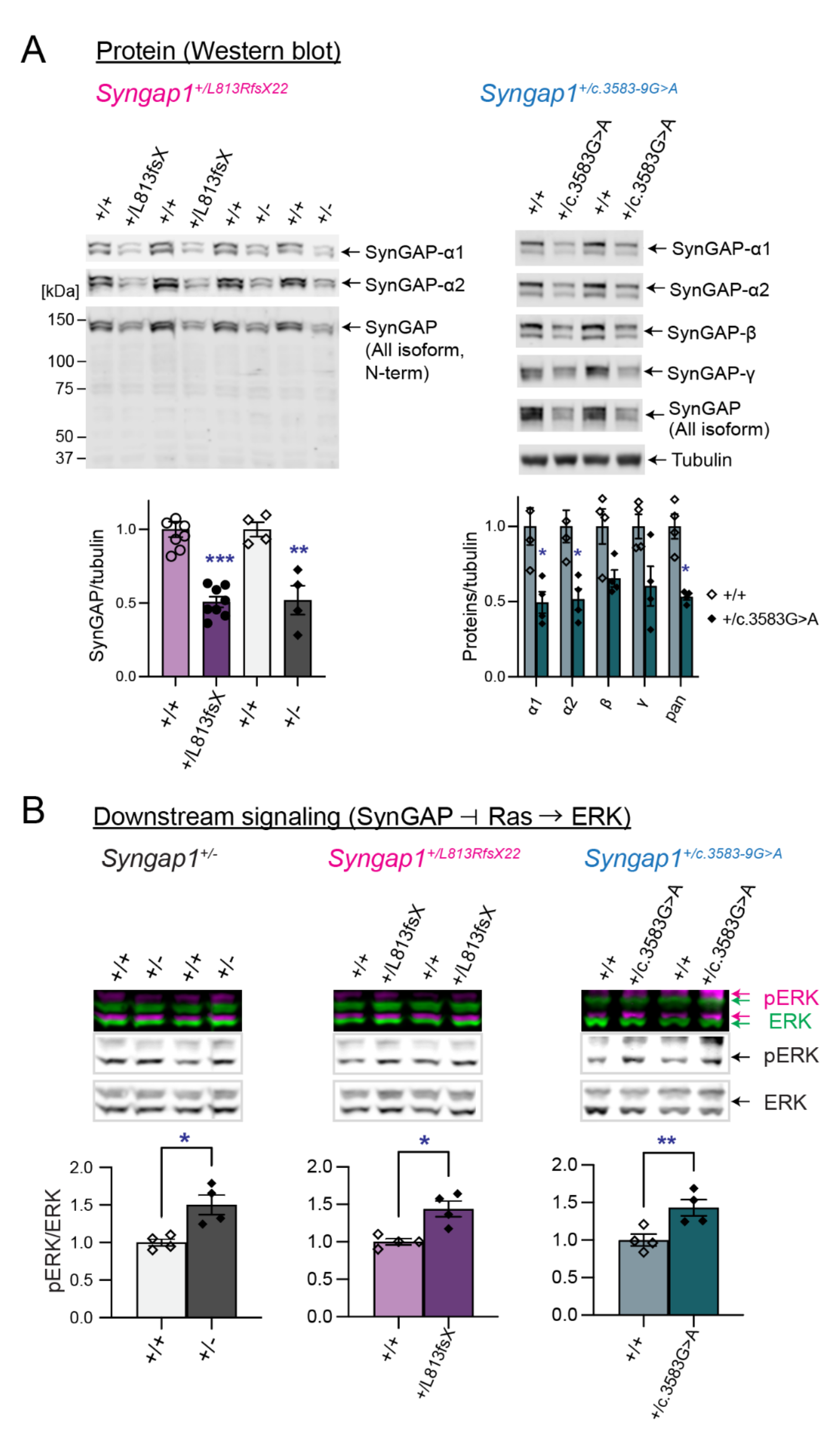
**Severe reduction of Syngap1 protein expression and aberrant downstream signaling (Syngap1-Ras-ERK) in SRID mice** (A) Western blotting of total Syngap1 proteins and their c-terminal isoforms. L813RfsX22; the western blotting using N-terminal antibody was conducted to check for any truncated protein expression. Truncated protein expression was not detectable. One-way ANOVA followed by Tukey’s post hoc test (Genotypes F (3,20) = 26.06; p < 0.0001, n = 4-7 independent brain samples, *** p<0.001, ** p<0.01, * < 0.05) was performed. c.3583-9G>A; Western blotting for all Syngap1 C-terminal isoforms is shown. Two-way ANOVA followed by Tukey’s post hoc test (Isoforms F (4,30) = 0.2621; p = 0.899, Genotype F (1,30) = 53.07; p < 0.0001, Interaction F (4,30) = 0.2621; p = 0.899, n = 3-4 individual brains, *** p<0.001, ** p<0.01, * < 0.05). (B) Western blots of phosphorylated ERK (active) / total ERK from Syngap1 mutant lines. ERK activation levels were quantified by calculating phosphorylated ERK levels normalized with total ERK levels. Two-tailed t-tests (* p< 0.05, ** p< 0.01, *** p<0.001).

### *Syngap1^+/c.3583-9G>A^*mice show 40-50% less *Syngap1* mRNA and Syngap1 protein

Total RNA was extracted from whole brains of *Syngap1^+/c.3583-9G>A^*mice and analyzed by northern blot and qPCR. Strikingly, brains from *Syngap1^+/c.3583-9G>A^* mice exhibited 40-50% less *Syngap1* mRNA by Northern blot (53.2 ± 3.3 %, p < 0.0001, **Fig. 2*A***) and qPCR (62.5 % ± 2.9 % compared to wild-type, p = 0.004, **Fig. 2*B***), indicating that the c.3583-9G>A mutation likely results in *Syngap1* mRNA undergoing NMD. Western blotting of protein extracted from whole brain of *Syngap1^+/c.3583-9G>A^* revealed approximately 50% less Syngap1 protein expression (53.2 ± 2.0, p = 0.0325, two-tailed t-test), similar to heterozygous *Syngap1* knock-out mice (*Syngap1^+/-^*). Protein expression of Syngap1 α1 and α2 C-terminal isoforms were reduced by approximately 50% (α1 49.5 ± 7.2 %, p = 0.016; α2 51.5 ± 7.0%, p = 0.024). There were trends toward decreased β and γ C-terminal isoform expression, but these were not statistically significant (β: 64.5% ± 1.7%, p= 0.234; γ: 60.3 ± 1.3%, p =0.114) (**Fig. 3*A***).

### Elevated Ras-ERK signaling in Syngap1 knock-out and SRID model mice

Syngap1 is a Ras-GTPase activating protein that negatively regulates Ras-ERK signaling in neurons (**Fig. 3*B***). To investigate ERK signaling in *Syngap1* mutant mice, protein was extracted from whole brain of heterozygous knock-out (*Syngap1^+/-^*) and SRID model mice, and western blotting was performed to quantify phosphorylated (active) ERK and total ERK. There was a large increase of phosphorylated ERK over total ERK that was comparable across all mutant mouse models (*Syngap1^+/-^*: 150.3 ± 13.0 %, p = 0.0425; *Syngap1^+/L813RfsX22^*: 144.0 ± 10.6 %, p= 0.027; *Syngap1^+/c.3583-9G>A:^* 143.0 ± 10.8 %, p= 0.018; two-tailed t-test).

### RNA-seq revealed aberrant splicing and converging downstream transcriptional changes in *Syngap1^+/c.3583-9G>A^ and Syngap1^+/-^* mice

To examine the impact of the cryptic splice site mutation in further detail and discover transcriptional changes due to *Syngap1* loss-of-function, we performed RNA-seq on whole-brain samples of *Syngap1^+/c.3583-9G>A^ and Syngap1^+/-^* mice. Splice junction reads confirmed that the predicted cryptic splice acceptor indeed led to aberrant splicing at the exon 16-17 junction of *Syngap1*, which was not observed in any of the other mouse models (**Fig. 4*A***). The large decrease (∼45%) in exon 16-17 junction coverage and the relative scarcity of the cryptic splicing junction suggests the transcripts undergo nonsense-mediated decay (NMD) due to the premature stop codon. The wild-type exon 17 displays a 6bp 5’ exon extension encoding the two amino acids valine and lysine (i.e., ‘VK’ exon extension). The ratio of VK vs non-VK splice junctions did not change in the *Syngap1^+/c.3583-9G>A^* mice, likely excluding the modulation of this splice decision as a factor for SRID in the person with this mutation (***SI Appendix*, Fig. S1*B***). Other alternative splicing events, including exon 14 skipping (36), exon 18 5’ extension (β vs. non-β), exon 18 3’ extensions and exon 19 skipping (α1, α2, γ) did not show significant changes in ratio (***SI Appendix*, Figs. S1*C*, D, E**). These results confirm the impact of the cryptic splice acceptor in *Syngap1^+/c.3583-9G>A^*mice is dominant to the wild-type acceptor and confined to the specific splicing site.

**Figure 4.**
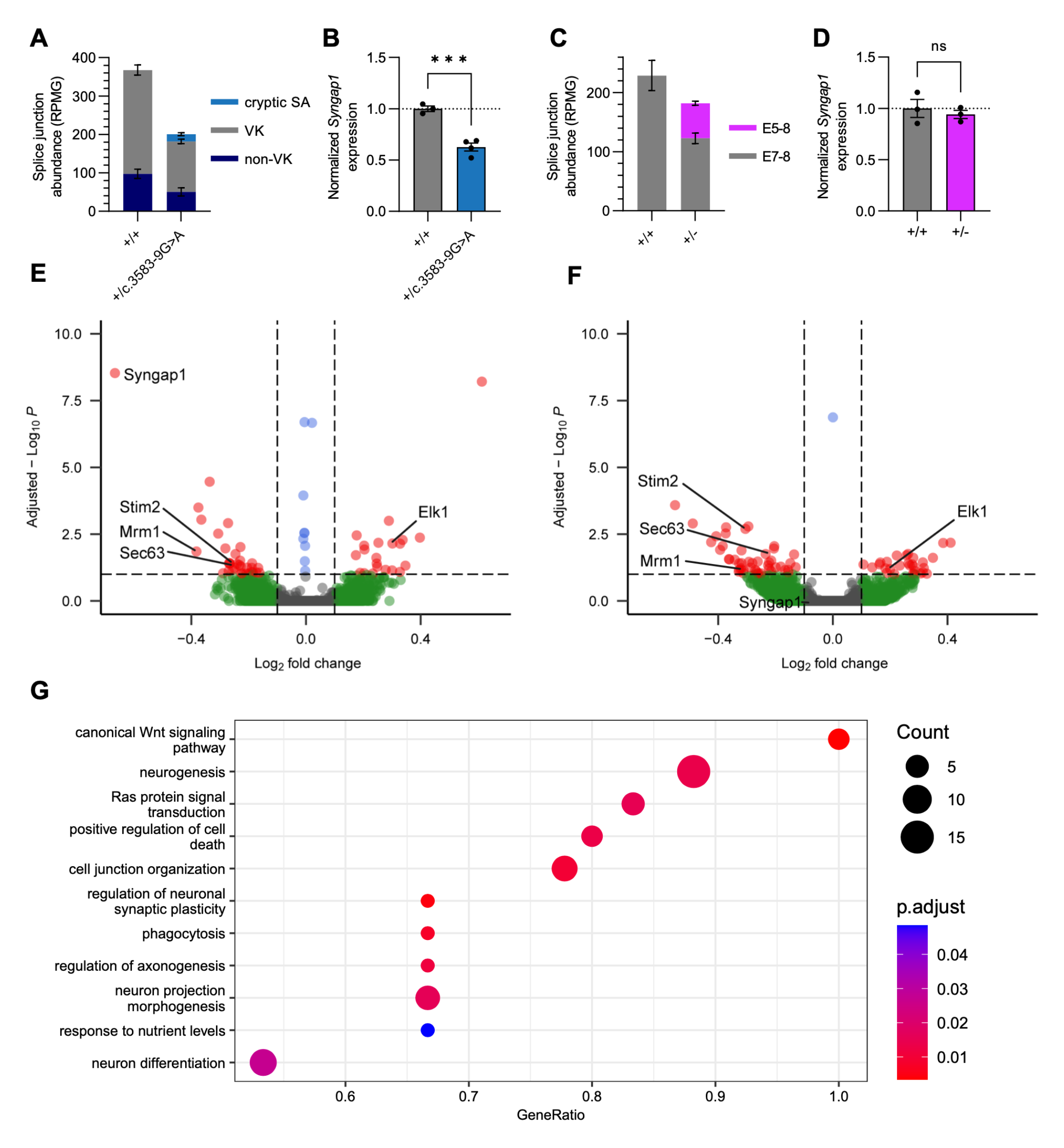
**RNA-seq reveals aberrant splicing and converging downstream transcriptional changes in *Syngap1^+/c.3583-9G>A^ and Syngap1^+/-^* mice** (A) Splice junction read abundance quantified as Reads Per Million Gapped (RPMG) between exons 16 and 17 of *Syngap1*. Reads with splice donor site at the 3’ end of exon 16 spliced to the ‘VK’ exon extension at the 5’ end of exon 17 and non-VK splice acceptor 6bp downstream in +/+ littermates, whereas only the Syngap1^+/c.3583-9G>A^ mice displayed a small fraction of cryptic splice acceptor site −7bp upstream, indicating efficient NMD. A two-way ANOVA revealed a significant effect of genotype, isoform, and interaction (Genotype F (1, 5) = 50.67; p = 0.0008, Isoform F (2, 10) = 297.7; p < 0.0001, Interaction F (2, 10) = 49.32; p < 0.0001, n = 3-4 samples/group) and Šídák’s post hoc tests confirmed significant decreases in VK (p < 0.0001) and non-VK splice junction abundance (p < 0.0044). (B) Total *Syngap1* expression was significantly decreased in Syngap1^+/c.3583-9G>A^ mice (+/+: 1.000 ± 0.027; +/c.3583- 9G>A: 0.626 ± 0.039; p=0.0007, unpaired t-test). (C) Splice junction read abundance in Syngap1^+/-^ mice. Reads with *Syngap1* splice acceptor site at the 5’ end of exon 8 spliced to the 5’ end of exon 7 in +/+ littermates, whereas only the Syngap1^+/-^ mice displayed aberrant exon 6 and 7 skipping, verifying the intended knockout of exons 6 and 7. Note the significant amount of exon 5-8 junction reads detected, which indicates inefficient NMD. A two-way ANOVA revealed a significant effect of isoform and interaction (Genotype F (1, 4) = 2.728; p = 0.1740, Isoform F (1, 4) = 124.7; p < 0.0004, Interaction F (1, 4) = 40.13; p < 0.0001, n = 3-4 samples/group) and Šídák’s post hoc tests confirmed significant decreases in exon 7-8 (p = 0.03) and exon 4-8 splice junction abundance (p < 0.0011). (D) Total *Syngap1* expression was not significantly decreased in Syngap1^+/-^ mice (+/+: 1.000 ± 0.087; +/-: 0.941 ± 0.040; p=0.5714, unpaired t-test). (E and F) Differential gene expression analysis of *Syngap1^+/c.3583-9G>A^*(E) and *Syngap1^+/-^* (F) mice reveals converging downstream regulated genes. (G) Gene set enrichment analysis of a merged dataset of both *Syngap1* loss-of-function mouse lines (*Syngap1^+/c.3583-9G>A^* and *Syngap1^+/-^*) revealed transcriptional regulation significantly enriched in a number of biological processes (BP). Gene ratio is the portion of genes that significantly are significantly regulated from the total number of genes associated to that process. Genes with increased and decreased expression were pooled to visualize general roles of *Syngap1*-regulated genes rather than the polarity of regulation in each pathway.

*Syngap1^+/-^* mice were previously generated by deleting the core exons 6 and 7, which shifts the reading frame and leads to reduction of Syngap1 (37). We verified in brain RNA-seq of these mice that exons 6 and 7 are skipped in a large portion of transcripts, which are not completely degraded through NMD (**Fig. 4*C***). Perhaps due to this incomplete NMD, total estimated *Syngap1* RNA expression is not significantly lower in these mice compared to wild-type littermates. The difference with qPCR and Northern blot quantification could be due to transcripts with intron retention and immature polyA tails, which affect these measures differently. Given the halved Syngap1 protein expression and absence of truncated protein species (**Figs. 2, 3*A***), this suggests that the transcripts from the knockout allele may be protected from NMD to an extent but are nevertheless translationally inactive. Substantial intron retention of intron 7 in *Syngap1^+/-^* mice (data not shown) further suggests nuclear retention of unspliced pre-mRNA may contribute to this resistance to NMD, which occurs in the cytosol (35). Alternative splicing events, including exon 14 skipping, exon 17 5’ extension (VK), exon 18 5’ extension (β vs. non-β), exon 18 3’ extensions, and exon 19 skipping (α1, α2, γ) did not show significant changes in ratio (***SI Appendix*, Figs. S1B-E**). Intriguingly, whereas an N-terminal splice junction unique to full length *Syngap1* (A1/2/4; exon 3-4) was significantly reduced in both *Syngap1^+/c.3583-9G>A^*(p = 0.0210, two-way ANOVA and Šídák’s post hoc test) and *Syngap1^+/-^* mice (p = 0.0437), other isoforms, including the second-most abundant D isoform, did not show a significant reduction (***SI Appendix*, Figs. S1F, G**). This suggests that these N-terminal isoforms may be subject to less efficient NMD or that they may be upregulated in a compensatory manner. The less pronounced down-regulation of these N-terminal isoforms might contribute to the <50% decrease observed in *Syngap1* mRNA levels in both heterozygous mouse lines. (**Figs. 2, 3*A***).

We next examined the transcriptome-wide changes that result from *Syngap1* loss-of-function in *Syngap1^+/c.3583-9G>A^ and Syngap1^+/-^* mice. Differential expression (DE) analysis revealed significant changes in gene expression: 54 (21 increased/33 decreased, ***SI Appendix*, Table S1**) and 76 (29 increased/47 decreased, ***SI Appendix*, Table S2**) in *Syngap1^+/c.3583-9G>A^ and Syngap1^+/-^* mice respectively, and demonstrated convergence with commonly regulated genes including decreased *Stim2*, *Mrm1*, *Sec63* and increased *Elk1* expression. Elk1 is a transcription factor involved in neuronal survival and plasticity that is phosphorylated by MAPK/ERK (38). Stim2 is crucial for maintaining calcium homeostasis in neurons, and plays a role in neuronal survival, synaptic plasticity, and memory (39).

The two mouse lines and littermate controls were respectively pooled for higher stringency and statistical power, which led to a refined list of 44 upregulated and 66 downregulated genes (***SI Appendix*, Table S3**). Gene Set Enrichment Analysis (GSEA) (40, 41) of these genes revealed strong enrichment of genes involved in the canonical Wnt signaling pathway (*Rps12, Nrarp, Aes, Sox2*), phagocytosis (*Pik3ca, Appl2*), regulation of neuronal synaptic plasticity (*Cntn2, Syngap1*), Ras protein signal transduction (*Plce1, Map4k4, Abl2, Syngap1, Gpsm2*), and neuron projection morphogenesis (*Map4k4, Sema4d, Abl2, Rab8a, Syngap1, Cntn2*), among others (**Fig. 4G**). Other top genes with significant changes in expression were *Wwox* (upregulated), a gene previously associated with NDD, *Kcnj9* (Kir3.3) (downregulated), which may account for intrinsic excitability changes, and *Camk4* (downregulated), a major downstream kinase of CaMKII essential for excitation-transcription coupling. These novel results demonstrate broad transcriptional changes in *Syngap1* mouse models and provide a foundation for further investigation of downstream pathways as well as for biomarker discovery.

### Mouse models of *SYNGAP1*-related intellectual disability exhibit synaptic plasticity deficits

In both mice and humans, loss of SYNGAP1 expression causes learning impairment that may result from deficits in synaptic plasticity (8, 9, 13, 24). We and others have demonstrated that heterozygous *Syngap1* knockout (*Syngap1^+/-^*) mice have deficits in theta-burst stimulated (TBS) induced long-term potentiation (TBS-LTP) of CA3 è CA1 synapses within the hippocampus (8, 9). Since *Syngap1^+/L813Rfsx22^*and *Syngap1^+/c.3583-9G>A^* mice exhibit an approximately 50% reduction in Syngap1 protein like *Syngap1^+/-^* mice, we next sought to determine if similar TBS-LTP deficits exist in each of the *Syngap1^+/L813Rfsx22^*and *Syngap1^+/c.3583-9G>A^* mouse models. Extracellular field recordings of TBS-LTP were performed in acute hippocampal slices of adult *Syngap1^+/L813Rfsx22^* and *Syngap1^+/c.3583-9G>A^* animals and compared to the magnitude of TBS-LTP measured in respective wild-type littermates (**Fig. 5**). Consistent with our data demonstrating that Syngap1 protein and mRNA level are reduced with single allelic mutation of either L813RfsX22 or c.3589G>A, TBS-LTP was similarly decreased by 45.2% and 50.1%, respectively (*Syngap1^+/L813RfsX22^*: 131.1 ± 5.80% LTP, n = 12; *Syngap1^+/+^*: 156.8 ± 7.88% LTP, n = 10; Mann-Whitney p = 0.0169)(*Syngap1^+/c.3583-9G>A^*: 134.9 ± 5.90% LTP, n = 14; *Syngap1^+/+^*: 169.9 ± 6.89% LTP, n = 18; Unpaired t-test, p = 0.0008). These data demonstrate that single allelic loss-of-function *Syngap1* mutations are sufficient to cause abnormal synaptic plasticity.

**Figure 5.**
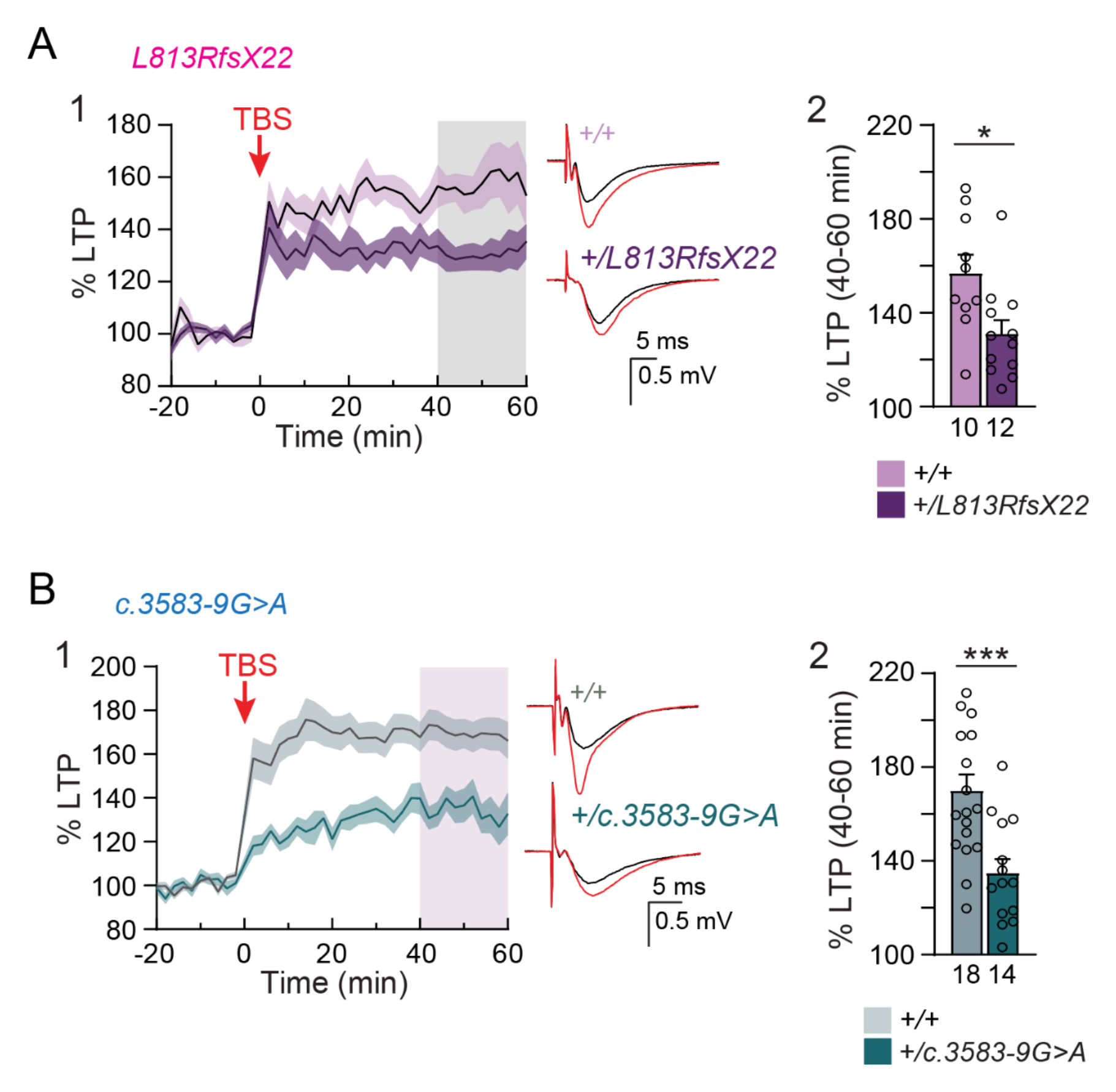
**SRID mice exhibit synaptic plasticity deficits** (A1) Averaged population field CA1 recordings of TBS-LTP time course obtained from brain slices of *Syngap1^+/L813RfsX22^* mice and wild-type (*Syngap1*^+/+^) littermates. All data points are normalized to the averaged baseline fEPSP slope. Inset: Example averaged fEPSP traces from *Syngap1*^+/+^ and *Syngap1^+L813RfsX22^* slices recorded during baseline (black) and 40-60 minutes after TBS-LTP induction (red). (A2) Quantification of averaged TBS-LTP in *Syngap1^+L813RfsX22^* and *Syngap1*^+/+^ littermates. Individual data points are superimposed. TBS-LTP is calculated by the ratio of the mean fEPSP slope measured 40-60 minutes after TBS-LTP induction (gray shaded region) divided by the averaged fEPSP baseline slope within each recorded sample. (*Syngap1*^+/+^: n = 10, 156.8 ± 7.88% S.E.M.; *Syngap1*^+/L813RfsX22^: n = 12, 131.1 ± 5.80% S.E.M.) Statistics: D’Agostino & Pearson test: non-normal distribution, Mann-Whitney ranked sum test, p = 0.0169. Error bars and shading represent the S.E.M. p < 0.05*, p < 0.01**, p < 0.001***. (B1) Averaged population field CA1 recordings of TBS-LTP time course obtained from brain slices of *Syngap1^+/c.3583-9G>A^*mice and *Syngap1*^+/+^ littermates. Inset: Example averaged fEPSP traces from *Syngap1^+/c.3583-9G>A^* and *Syngap1*^+/+^ slices recorded during baseline (black) and 40-60 minutes after TBS-LTP induction (red). (B2) Quantification of averaged TBS-LTP in *Syngap1^+/c.3583-9G>A^*mice and *Syngap1*^+/+^ littermates. Individual data points are superimposed. (*Syngap1*^+/+^: n = 18, 169.9 ± 6.89% S.E.M.; *Syngap1^+/c.3583-9G>A^*: n = 14, 134.9 ± 5.90% S.E.M.) Statistics: D’Agostino & Pearson test: normal distribution, Unpaired t-test, p = 0.0008. Error bars and shading represent the S.E.M.. p < 0.05*, p < 0.01**, p < 0.001***.

### *Syngap1* mutant mice show hyperactivity and working memory impairment

To test working memory, we performed spontaneous Y maze behavioral testing in both mutant mouse models (**Fig. 6*A***). Compared to wild-type mice, heterozygous *Syngap1^+/L813Rfsx22^* mice demonstrated decreased spontaneous alternations (*Syngap1^+/+^* 72.3 ± 2.5 % alternation, *Syngap1^+/L813Rfsx22^* 54.4 ± 2.3 % alternation, p < 0.001 *** Two-tailed T-test) (**Fig. 6*B,*** Left panel), similar to heterozygous *Syngap1* knock-out mice, *Syngap1*^+/-^ (13). Both *Syngap1^+/L813Rfsx22^* and *Syngap1*^+/-^ mice also showed an increased number of repetitive arm visits (*Syngap1^+/+^* 6.0 ± 0.9 repetitions, *Syngap1^+/L813Rfsx22^* 15.6 ± 1.4 repetitions, p < 0.001 *** Two-tailed T-test) (**Fig. 6*B,*** Middle panel), a measure of repetitive behavior, which is a known clinical finding in SRID (24, 28). The number of arm entries over 5 minutes was also significantly increased in *Syngap1^+/L813Rfsx22^* (*Syngap1^+/+^* 21.5 ± 2.6 entries, *Syngap1^+/L813Rfsx22^* 33.3 ± 2.7 entries, p < 0.01 ** Two-tailed T-test) (**Fig. 6*B,*** Middle panel), similar to *Syngap1*^+/-^ mice (13).

**Figure 6.**
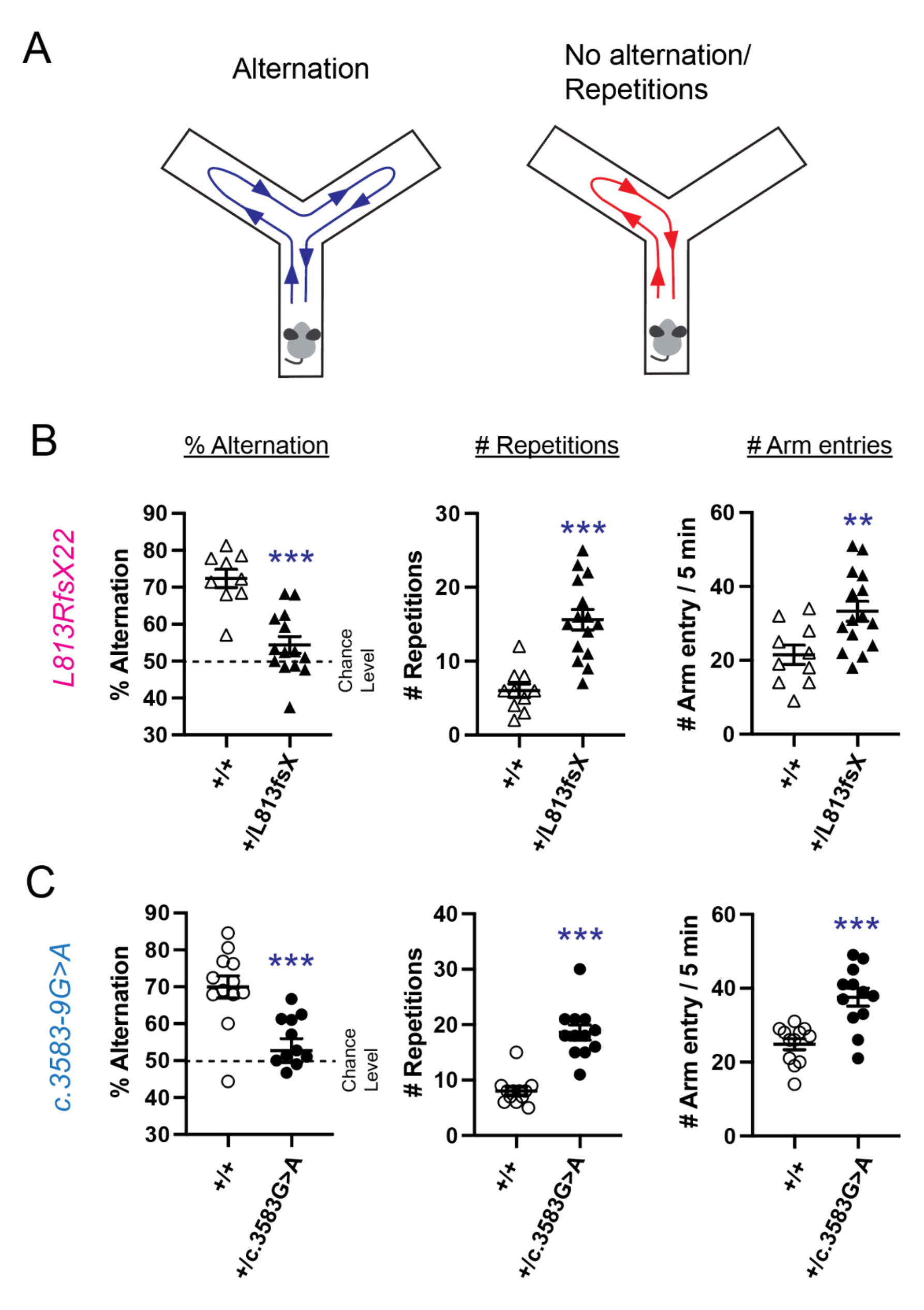
**Recapitulation of working memory deficits, repetitive behavior, and hyperactivity in SRID model mice** (A) Diagram of the experimental Y-maze setup. An arm entry was recorded as an alternation when the mouse fully entered an arm that it had not visited most recently (e.g., Arm A to Arm B to Arm C is an alternation; Arm A to Arm B to Arm A is not an alternation). (B) Spontaneous alternation rate of arm visits (% alternation), the number of repetitive arm visits (# repetitions), and the number of arm visits (# arm entries) for wild-type (*Syngap1^+/+^*) or *Syngap1*^+/L813RfsX22^ are shown. Two-tailed t-test was performed (* p< 0.05, ** p< 0.01, *** p<0.001). (C) % alternation, # repetitions, and # arm entries of for wild-type (*Syngap1^+/+^*) or *Syngap1^+/c.3583-9G>A^*are shown. Two-tailed t-test was performed (* p< 0.05, ** p< 0.01, *** p<0.001).

These changes were observed in *Syngap1^+/L813Rfsx22^* mice of both sexes including alternations (Male: *Syngap1^+/+^* 73.6 ± 2.4 % alternation, *Syngap1^+/L813Rfsx22^* 52.7 ± 3.2 % alternation, p < 0.05 * : Female; *Syngap1^+/+^* 71.5 ± 3.8 % alternation, *Syngap1^+/L813Rfsx22^* 52.3 ± 5.3 % alternation, p < 0.05 *; One-way ANOVA followed by Tukey test) (***SI Appendix*, Figs. S2*A,*** Left panel), repetitive arm visits (Male: *Syngap1^+/+^* 5.8 ± 1.3 repetitions, *Syngap1^+/L813Rfsx22^* 15.6 ± 1.8 repetitions, p < 0.05 * : Female; *Syngap1^+/+^* 6.2 ± 1.3 repetitions, *Syngap1^+/L813Rfsx22^* 15.7 ± 2.5 repetitions, p < 0.05 *; One-way ANOVA followed by Tukey test) (***SI Appendix*, Figs. S2*A,*** Middle panel) and number of arm entries (Male: *Syngap1^+/+^* 21.7 ± 4.8 entries, *Syngap1^+/L813Rfsx22^* 33.3 ± 3.7 entries, p = 0.25 : Female; *Syngap1^+/+^* 21.3 ± 3.3 entries, *Syngap1^+/L813Rfsx22^* 33.1 ± 4.3 entries, p = 0.20; One-way ANOVA followed by Tukey test) (***SI Appendix*, Figs. S2*A,*** Right panel).

Next, we conducted the spontaneous alternation Y-maze testing in *Syngap1^+/c.3583-9G>A^* mice (**Fig. 6*C***). Similar to *Syngap1*^+/-^ mice and *Syngap1^+/L813Rfsx22^*mice, *Syngap1^+/c.3583-9G>A^* mice showed a decreased number of spontaneous alternations (*Syngap1^+/+^* 69.9 ± 2.9 % alternation, *Syngap1^+/c.3583-9G>A^* 52.7 ± 3.2 % alternation, p < 0.001 *** Two-tailed T-test; **Fig. 6*C,*** Left panel), more repetitive arm entries (*Syngap1^+/+^* 8.0 ± 0.8 repetitions, *Syngap1^+/c.3583-9G>A^*18.6 ± 1.3 repetitions, p < 0.001 *** Two-tailed T-test; **Fig. 6*C,*** Middle panel), and an increased number of total arm entries in 5 minutes (*Syngap1^+/+^*24.8 ± 1.5 entries, *Syngap1^+/c.3583-9G>A^* 37.6 ± 2.4 entries, p < 0.001 *** Two-tailed T-test; **Fig. 6*C,*** Middle panel).

Again these findings were observed in both sexes including decreased alternations (Male; *Syngap1*^+/+^ 74.5 ± 6.2 % alternation, *Syngap1^+/c.3583-9G>A^* 57.6 ± 3.1 % alternation, p < 0.05 *: Female; *Syngap1*^+/+^ 63.5 ± 5.4 % alternation, *Syngap1^+/c.3583-9G>A^* 47.9 ± 5.3 % alternation, p = 0.07; One-way ANOVA followed by Tukey test; ***SI Appendix*, Figs. S2*B,*** Left panel), increased repetitive arm visits (Male; *Syngap1*^+/+^ 7.5 ± 0.4 repetitions, *Syngap1^+/c.3583-9G>A^* 16.3 ± 1.4 repetitions, p < 0.01 ** : Female; *Syngap1*^+/+^ 8.6 ± 1.7 repetitions, *Syngap1^+/c.3583-9G>A^* 20.8 ± 2.0 repetitions, p < 0.001 ***; One-way ANOVA followed by Tukey test; ***SI Appendix*, Figs. S2*B,*** Middle panel), and an increased number of arm entries in 5 minutes (Male; *Syngap1*^+/+^ 25.4 ± 2.3 entries, *Syngap1^+/c.3583-9G>A^* 39.5 ± 4.4 entries, p < 0.05 * : Female; *Syngap1*^+/+^ 24.0 ± 1.9 entries, *Syngap1^+/c.3583-9G>A^* 35.6 ± 2.3 entries, p = 0.06; One-way ANOVA followed by Tukey test; ***SI Appendix*, Figs. S2*B,*** Right panel).

## Discussion

This is the first investigation of knock-in mouse models with pathogenic *Syngap1* variants found in people with SRID. Using CRISPR-Cas9 technology, we generated one SRID mouse model harboring a frameshift mutation, L813RfsX22, and a second model with an intronic mutation that creates a cryptic splice site, c.3583-9G>A. While *Syngap1* transcript levels are decreased by ∼35-40% in these models, importantly both models show reduction of Syngap1 protein by half, deficits in synaptic plasticity, and abnormal behavior including hyperactivity, repetitive behavior, and impaired working memory. The direct causality and severity of disease-associated human mutations (especially with splicing-related mutations) are difficult to predict without generating an animal model, and our results establish both mutations as equally sufficient to cause SRID phenotypes.

### Model mice recapitulate key endophenotypes of SRID

The clinical features of SRID include global developmental delay, seizures, skeletal abnormalities, hypotonia, strabismus, constipation, failure to thrive, hyperactivity, autistic behaviors including repetitive behavior and social deficits. Prevalence of SRID in males and females is approximately equal and there is no known sexual predominance (29, 42). Like humans with SRID, both *Syngap1^+/L813Rfsx22^* and *Syngap1^+/c.3583-9G>A^*mice display working memory impairment, hyperactivity, and repetitive behavior. Importantly, these findings were observed irrespective of sex (***SI Appendix*, Figs. S2 *A*, *B***), which is consistent with human data on cognitive function in males and females with SRID (28). Previously, several groups have found that many behavioral phenotypes are shared between male and female *Syngap1*^+/-^ mice (43-46). In contrast, one group reported decreased latency to fall in the rotarod test only in females (47) and another study showed a correlation in PSDs between decreased steady-state Syngap1 protein and higher amounts of TARPs only in females and not in males (48). Further studies are necessary to clarify the effect of sex on the phenotypes of the *Syngap1*^+/-^ as well as the novel SRID mouse lines used in the present study. Our findings provide two mouse models that recapitulate behaviors found in SRID in both sexes and will be valuable resources to further study features of SRID including epilepsy and abnormal social interaction.

### Findings in SRID mice support a loss-of-function mechanism for disease pathogenesis

Supporting a loss-of-function mechanism of disease, both SRID mouse models show a reduction in *Syngap1* mRNA that varies from 30-50% depending on the specific mutation. With Northern blots, qRT-PCR, and RNA-seq, we confirmed the impact of the cryptic splice acceptor in *Syngap1^+/c.3583-9G>A^* mice is dominant to the wild-type acceptor and is confined to the specific splicing site. This is important due to the complex nature of splicing regulation and is relevant for therapeutic development. We also found that transcripts lacking exons 6 and 7 in *Syngap1^+/-^*mice can persist in a presumed translationally inactive state, as do some N-terminal isoforms including the D isoform. The mechanism of such escape from NMD is not clear.

Both SRID mouse models also show reduced Syngap1 protein and behaviors that resemble those in SRID. While homozygous knockout (*Syngap1*^-/-^) mice die within a week of birth, previous studies have shown heterozygous knock-out mice (*Syngap1*^+/-^) are viable and show increased RAS/MAPK signaling and long-term potentiation impairment at Shaffer collateral-CA1 synapses in the hippocampal slices, as well as seizures and behavior abnormalities including hyperactivity, social deficits, and poor working memory (43, 44, 46, 47, 49, 50). As behavioral deficits in *Syngap1*^+/-^ mice closely resemble those found in SRID, it has been hypothesized that that a loss-of-function mechanism underlies SRID pathogenesis (13). Here we show that two different knock-in mouse models with known SRID mutations indeed both show half the normal amount of Syngap1 protein and phenotypically recapitulate multiple clinical features of SRID, implicating NMD and *SYNGAP1* haploinsufficiency as the core of pathogenesis.

### RNA-seq of *Syngap1* mice enables transcriptome-wide discovery of downstream changes and potential biomarkers

Our RNA-seq data revealed highly significant changes in several genes associated with synaptic plasticity, intrinsic excitability, transcription factors, and NDD including *Stim2*, *Elk1, Aes* (*Tle5*), *Mrm1*, *Sec63, Wwox*, and *Camk4*, suggesting widespread transcriptional changes that may contribute or counteract the phenotypes of SRID. These novel results will aid the characterization of SRID pathophysiology and provide candidate biomarkers for diagnosis and treatment.

### New *Syngap1* mouse models for SRID treatment development

The present study shows that mice with distinct SRID mutations recapitulate phenotypic features of SRID and provides a framework for new areas of therapeutic intervention. While case reports suggest potential efficacy of various medications including statins (51), more research is needed and currently there is no standard-of-care disease-modifying treatment for SRID (28). In preclinical literature, interventions including lovastatin treatment in hippocampal slices (52) and acute perampanel treatment (52) have been explored. However, studies showing definitive phenotypic rescue in *Syngap1*^+/-^ mice with pharmacologic treatment are lacking. For example, treatment of *Syngap1*^+/-^ mice with the MEK inhibitor PD-0325901 did not improve LTP impairment (53). Our findings establish two novel mouse lines as excellent models to further interrogate SRID pathophysiology and test potential treatments. These mouse models will complement existing *Syngap1* mutant mice and other model animals by providing a diversity of causal mutations, to accelerate safe and generally applicable therapeutic development.

## Materials and Methods

### Reagents

All restriction enzymes were obtained from New England Biolabs. Chemicals were obtained from SIGMA-Aldrich unless otherwise specified. TTX, Bicuculline, and Strychnine were obtained from TOCRIS Bioscience. Goat anti-SynGAP-α1 antibody is from Santa Cruz (sc-8572). Rabbit pan-SynGAP 947-1167 antibody is from Thermo scientific (#PA-1-046). DNA sequencing was performed at the Johns Hopkins University School of Medicine Sequencing Facility.

### Animals

*Syngap1^+/c.3583-9G>A^ mutants, Syngap1^+/L813RfsX22^*mutants, and wild-type littermates were maintained on a mixed background of C57/B6J and 129/SvEv background strains. All animals were housed in the Johns Hopkins University animal facility. Animals were allowed *ad libitum* access to food and water and were reared on a typical 12-hour light-dark cycle. All animal experiments utilized both male and female mice at specified ages and were conducted in accordance with the guidelines implemented by the Institutional Animal Care and Use Committee at Johns Hopkins University.

### CRISPR/Cas9-based mouse gene engineering

All mouse gene engineering steps were performed by the Johns Hopkins Transgenic Core. One-cell stage fertilized C57BL/6 mouse embryos were injected by Cas9 protein, crRNA, tracrRNA, and homology directed repair DNA template. Guide RNA sequences are 5’- GAGCTGCTCGTGCAGTATGC-3’ for L813RfsX22 and 5’-GTACGGGGTCATGTGCCCGG-3’ for c.3583-9G>A. Homology directed repair donor templates are 5’- ACCACCACCCGGTGGGGGTAAAGACCTTTTCTATGTGTCTAGACCTCCTCGACCAGTTCCACATCAACATACTGCACGAGCAGCTCGGACATCACAGAGCCAGAGCA-3’ (plus strand) for L813RfsX22 and 5’-CGAATGTATCTCCTCCTCATACTCCTTCACCTGCCAGCTGGGCACACGACCCCGCACTATGAGGGGCGCCCAGCCTTGGCTTTACCAGCCCACTCCCAT-3’ (minus strand) for c.3583-9G>A. Donor templates and crRNAs were synthesized by IDT. Offspring were screened by PCR with primers flanking the introduced mutations followed by diagnostic restriction digests. For L813RfsX22 forward (5’- TTGCTTCCAACAGCTCTATGGAC −3’) and reverse (5’- AACACTGCTACTGTTAAGGCGAC −3’) primers amplify a 256 bp PCR product. After digestion with XbaI mutant products will be cut to generate 154 and 102 bp fragments. For c.3583-9G>A forward (5’- ACCACCTTGAAGAAGCCTCAG −3’) and reverse (5’- GCAACCTCCGCTCATACTCT −3’) primers amplify a 270 bp product. After digestion with PvuII mutant products will be cut to generate 180 and 90 bp fragments. Sanger sequencing was performed on all mutant mice to confirm that HDR donor templates were accurately introduced into the genome.

### Southern Blotting

The overall structure of genome before and after recombination was confirmed by Southern blotting using standard techniques as previously described (8). For the L813RfsX22 mutation, a 400 bp DNA fragment upstream of exon15 was used as probe. For c.3583-9G> a, a 445 bp DNA fragment upstream of exon 17 was used as probe. Probes were labeled with ^32^P using the Prime-It II Random Primer Labeling Kit (Agilent Cat. #300385).

### RNA extraction and Northern blotting

Total brain RNA was isolated by TRIzol (Invitrogen Cat. #15596026). Samples were homogenized in TRIzol. Chloroform was added to homogenates and the samples were shaken vigorously for 15 s. Samples were incubated at room temperature for 3 min and centrifuged at 13,000 × g for 15 min at 4 °C. The aqueous phase was carefully removed and applied to a genomic DNA elimination column (approximately 350 µl) (Qiagen RNeasy Plus kit, catalogue no. 74136). The column was centrifuged for 30 s at 13,000 × g. After extraction, RNA concentration was measured using a Nanodrop (Thermo Scientific) and stored at –80 °C. 10 μg of total RNA was subjected to electrophoresis in a 0.9% denaturing agarose gel submerged in MOPS buffer (20mM MOPS, 5mM Sodium Acetate, 1mM EDTA, pH 7.0). RNA was transferred to Hybond-N+ membranes (GE Healthcare, Cat #RPN303B) by the capillary transfer method using blotting paper. cDNA probes were labeled with ^32^P using the Prime-It II Random Primer Labeling Kit (Agilent Cat. #300385). The *Syngap1* probe corresponds to NM_001281491 nucleotides 1361- 2002. The *Rps26* probe corresponds to NM_013765.2 nucleotides 44-381. Membranes were hybridized overnight at 65°C with probes in SDS-PIPES buffer (50 mM PIPES, 100 mM NaCl, 50 mM NaH_2_PO_4_, 1 mM EDTA, 5% SDS pH6.8), washed and visualized by autoradiograms.

### Quantitative Reverse Transcription PCR (qRT-PCR)

500 ng RNA from each sample was reverse-transcribed with Superscript IV (Invitrogen). qPCR amplifications were carried out in 96-well plates using a CFX Connect (Bio-Rad). The following TaqMan assay was used for *Syngap1*: Forward Primer 5’-CCGGACCAGCAGCTTTC Reverse Primer 5’-CCCAGGATGGAGCTGTG, Probe 5’-CCGAAGTGCTGACCATGACCGG. mRNA expression levels were normalized to the housekeeping gene *Actb*, using the Mm.PT.58.29001744.g (IDT) assay in a multiplexed fashion. Results were calculated with the 2^-ΔΔCt^ method.

### RNA-seq library preparation and analysis

RNA samples were enriched for mRNA through bead-based polyA selection and libraries were generated with the NEBNext Ultra RNA Library Prep Kit (Illumina). cDNA libraries were barcoded and sequenced together on an Illumina Hiseq 4000 sequencer, generating 2×150-bp paired-end (PE) reads. The sequencing library was validated on the Agilent TapeStation (Agilent Technologies), and quantified by using Qubit 2.0 Fluorometer (Invitrogen) as well as by quantitative PCR (KAPA Biosystems).

The RNA-Seq pipeline from the bcbio-nextgen project (https://doi.org/10.5281/zenodo.3564938) and the bcbioRNASeq R package (54) were used to process and analyze all samples. The alignment of reads to the Genome Reference Consortium Mouse Reference build number 38 (GRCm38) of the mouse genome (mm10), which was supplemented with transcript information from Ensembl, was performed using STAR (55). FeatureCounts (56) was used to generate counts of reads aligning to known genes, which were then used in quality control measures. Gene counts were computed with the fast inference algorithm of Salmon (57) and imported with tximport. The quality of STAR alignments was assessed for evenness of coverage, ribosomal RNA content, exon and intron mapping rate, complexity, and other criteria using FastQC (https://www.bioinformatics.babraham.ac.uk/projects/fastqc/), Qualimap (58), and MultiQC (59). Both principal component analysis and hierarchical clustering methods were used to cluster samples in an unsupervised manner. This was done using rlog transformed reads to identify possible outliers and technical artifacts. Samples exhibiting low mapping rates (<70%) or low RIN values and 5’>3’ biases were excluded from the analysis.

Differential expression at the gene level was determined using DESeq2 (60) with a false discovery rate of 0.1 and absolute log2 fold change value threshold of 0.1, correcting for rRNA ratio and sex. Genes with a base mean value of less than 100 were discarded. Gene set enrichment analyses (GSEA) were performed on lists of differentially expressed genes (DEGs) for GO biological pathway (BP) term enrichment without cut-offs using clusterProfiler (41) and fold change calculations from DESeq2. Functional gene sets with a false discovery rate-adjusted p value less than 0.05 were considered enriched. spliceSites (https://github.com/wokai/spliceSites) was used to quantify splice donor and acceptor sites. For N-terminal isoforms, we follow the nomenclature previously proposed (21). We follow the exon numbering of full-length mouse *Syngap1* A2-γ (gamma) transcript (XM_006524243.2), with the last exon of α1/α2 designated as exon 20.

### Western blotting

Brain tissue was excised from C57BL6 mice at specified ages. Tissue was lysed in 10 volumes of lysis buffer (50 mM Tris pH 8.0, 100 mM NaCl, 1 mM EDTA, 1 mM EGTA, 1% Triton X-100, 0.2% SDS, 0.5 % Sodium deoxycholate, with cOmplete Protease inhibitor EDTA-free mix (Roche/ SIGMA) by Dounce A homogenizer. Protein concentrations were measured by Pierce BCA assay kit (Pierce 23225). Equal protein amounts (10 μg) were loaded into each lane. After probing by primary and secondary antibodies, signals were measured by a fluorescence-based imaging system for our quantitative western blotting (Odyssey® CLx Imaging System). Fluorescence detection is suitable for quantitative immunoblotting across large dynamic ranges (61-64). 50% of the first experimental lane was run in the left-most lane in order to assure the given quantification is linear in every primary-secondary antibody combination.

### Electrophysiology Acute slice preparation

*Syngap1^+/c.3583-9G>A^* mice and *Syngap1^+/L813RfsX22^*mice (4-7 months of age), along with their respective wild-type littermates, were transcardially perfused with ice cold oxygenated (95% O_2_/5% CO_2_) dissection buffer (212.7 mM sucrose, 5 mM KCl, 1.25 mM Na_2_PO_4_, 10 mM glucose, 26 mM NaHCO_3_, 0.5 mM CaCl_2_, 10 mM MgCl_2_) under isoflurane anesthesia immediately prior to decapitation. The brain was rapidly removed from the skull, and the anterior surface of the brain was cut at ∼15 deg with respect to the anatomical coronal plane with the cut penetrating deeper along the ventral-to-dorsal axis in continuously oxygenated dissection buffer. Acute transverse hippocampal slices (400 µm thickness) were then prepared using a vibratome (Leica VT1200S) and were briefly washed of the sucrose-based dissection buffer in oxygenated artificial cerebrospinal fluid (ACSF) composed of (in mM): 119 NaCl, 5 KCl, 1.25 Na_2_PO_4_, 26 NaHCO_3_, 10 glucose, 2.5 CaCl_2_, and 1.5 MgCl_2_. Slices recovered in a chamber containing ACSF at 30°C for 30 minutes and then transferred to room temperature for an additional 60 minutes or until used for electrophysiological recordings. The experimenter was blinded to genotype until all experiments and analyses were completed.

### Extracellular LTP recordings

Slices were placed in a submersion recording chamber with recirculating oxygenated (95% O_2_/5% CO_2_) ACSF at 30°C. Synaptic field excitatory postsynaptic potentials (fEPSPs) were evoked in response to electrical stimulation of the Schaeffer collateral inputs via a bipolar theta glass Ag/AgCl electrode (3 MΩ) containing ACSF. The baseline stimulation intensity was determined prior to recording by measuring the stimulation that is sufficient to evoke a half maximal fEPSP amplitude, which is half of the threshold for eliciting a population spike. Upon starting an LTP recording, the baseline stimulation intensity was used to measure the fEPSP slope over a stable 20-minute baseline period in response to a single 0.2-ms stimulation pulse delivered every 30 seconds. Absolute inclusion criteria for sample LTP recordings required a minimum stable baseline period of 10 minutes whereby the baseline fEPSP slope did not drift by >10%. To induce LTP, 4 episodes of theta-burst stimulation (TBS) were triggered at 0.1 Hz. Each TBS episode consisted of 10 stimulus trains administered at 5 Hz, whereby one train consists of 4 pulses at 100 Hz. Following TBS, fEPSP slope was measured for an additional 60 minutes by delivering single electrical pulses every 30 seconds. The magnitude of LTP was quantified by normalizing the fEPSP slope to the average baseline response, and then calculating the average fEPSP slope between 40-60 min after TBS. Recordings were first performed in *Syngap1^+/L813RfsX22^* mice and wild-type littermates in an alternating fashion across experimental days until data acquisition was complete to best control for day-to-day experimental variability. Subsequently, the same alternating recording pattern was implemented during LTP recording data acquisition from *Syngap1^+/c.3589GèA^* and wild-type littermates. Statistical comparisons were made exclusively between either *Syngap1^+/c.3589GèA^* mice or *Syngap1^+/L813RfsX22^*mice and wild-type littermates. with a student t-test or Mann-Whitney test. p<0.05*,0.01**, p<0.001***.

### Behavior: Spontaneous alternation in Y-maze

Mice aged 4-6 months were subjected to the Y-maze spontaneous alternation task in order to assess working memory performance. All groups were approximately evenly divided (45-55%) between males and females.

#### Y-maze spontaneous alternation

Following a 30-minute acclimatization period, mice were placed in the center of a three-chamber Y-maze in which the three arms were oriented 120 degrees from one another. Mice were allowed to explore the apparatus for 5 minutes. Arm entries were recorded when both rear paws passed over the boundary line between the center region and arm region of the apparatus. An arm entry was recorded as an alternation when the mouse fully entered an arm that it had not visited most recently (e.g., Arm A to Arm B to Arm C is an alternation; Arm A to Arm B to Arm A is not an alternation). Percent alternation was calculated as the number of alternating arm entries divided by the total number of arm entries. The Y-maze apparatus was thoroughly cleaned between trials. Arm entries were recorded manually. The experimenter was blinded to genotype until all experiments and analyses were completed.

### Statistics

All data are expressed as means ± S.E.M. of values. Data distributions were tested for normality using specified methods. Parametric tests were used if the data were normally distributed, and non-parametric tests were otherwise used, as detailed in the text. For parametric tests, unpaired/paired t-tests, 1-way/2-way ANOVA tests with Tukey’s or Šídák’s post-hoc multiple comparison correction were used. For data that did not follow normal or log-normal distributions, the Mann–Whitney U-test and Kruskal-Wallis 1-way ANOVA test were used where appropriate. If a significant interaction between two-factors was observed by two-way ANOVA, multiple comparison-corrected Tukey *post hoc* tests were performed to compare the measures as a function of one factor in each fixed levels of another factor unless otherwise specified. Statistical analyses and preparations of graphs were performed using Excel 16, or GraphPad Prism 9.0 software (\**p* < 0.05; \*\**p* < 0.01; \*\*\**p* < 0.001).

## Acknowledgements

We thank all members of the Huganir lab for discussion and support throughout this work especially Shaowen Ju, Ria Oba, Yinuo Han, Hai Tran, August Tingjiao Li, Sam Myung, Sang Ho Kwon, Stephanie Glavaris, Manisha Pradhan, Drs. Timothy R. Gamache, Mengnan Tian, and Bian Liu for reagent preparation and critical reading of the manuscript. This work was supported by grants from the National Institute of Health (MH112151, NS036715), the SynGAP Research Fund (2020-2021), and the Simons Foundation (SFARI Pilot Award 731581). We want to thank the SynGAP Research Fund, the SYNGAP1 Foundation, and individuals with SRID. and their families for their support and advocacy.

