## Supplemental Material for "Mouse models of *SYNGAP1*-related intellectual disability"

**This PDF file includes:**

Figures S1 to S2
Tables S1 to S3

Supplementary Figure. 1

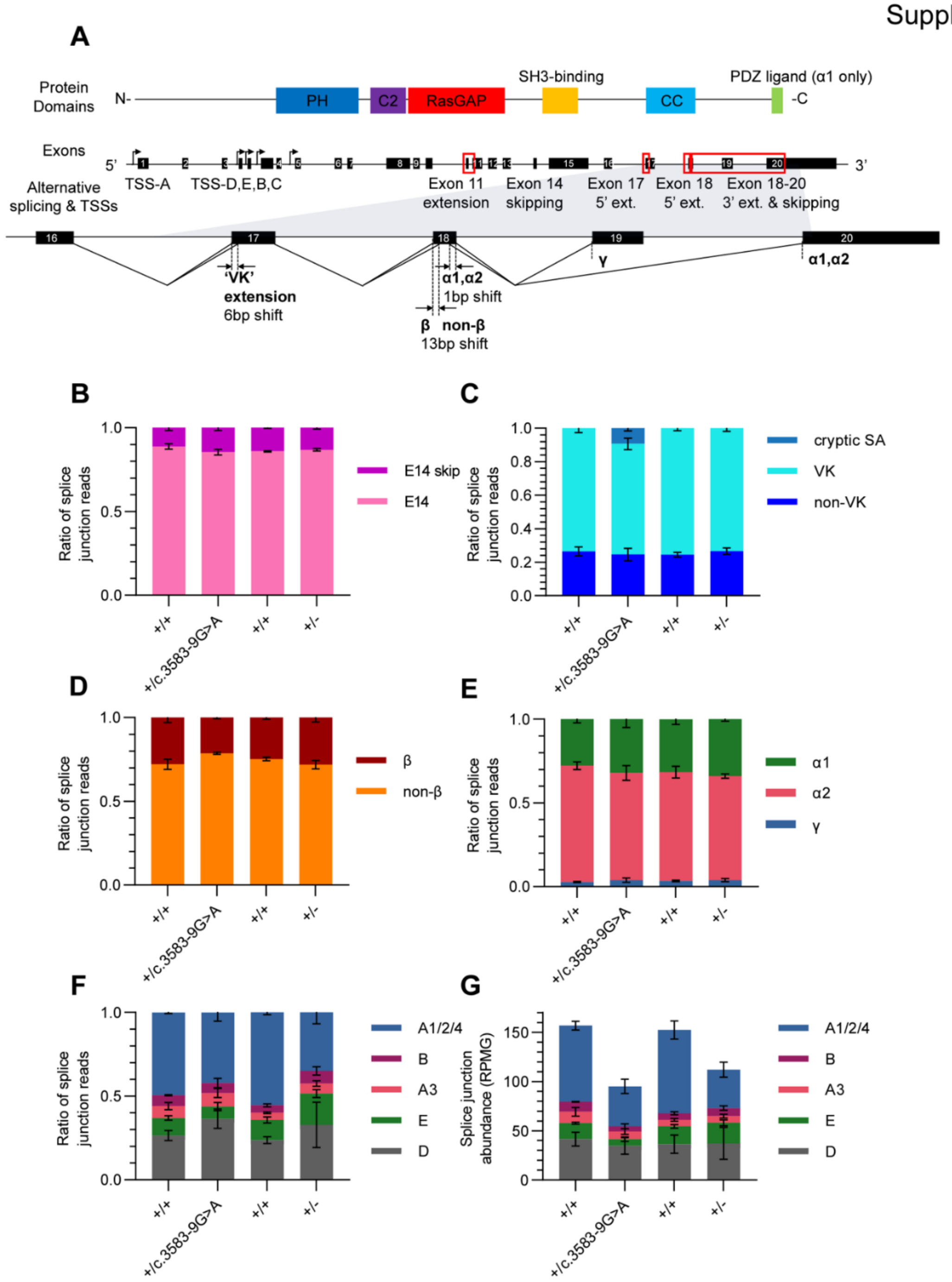


**Figure S1. Expression of alternative splice junctions in in *Syngap1*$^{+/c.3583\text{-}9G>A}$ and *Syngap1*$^{+/-}$ mice**

(A) Overview of gene structure of mouse *Syngap1*. Major protein domains, exons, alternative splicing events, and transcription start sites (TSS) are shown. The inset below shows a magnified view of the c-terminal exons 16-20.

(B) Exon 14 skipping ratio quantified as portion of junction reads (RPMG) splicing to exon 15 of *Syngap1*. Reads with the *Syngap1* exon 15 splice acceptor site at the 5' end of exon 15 spliced from the 3' end of exon 13 and 14 in all mice at similar ratios. This exon 14 skipping is more abundant outside the brain (data not shown). A two-way ANOVA revealed a significant effect of isoform only (Genotype $F (3, 10) = 0.000$; $p > 0.9999$, Isoform $F (1, 10) = 2730$; $p < 0.0001$,

Interaction $F (3, 10) = 1.285$; $p = 0.3324$, $n = 3\text{-}4$ samples/group) and Šídák's post hoc tests confirmed no significant differences between groups.
(C) Exon 17 VK/non-VK alternative splicing ratio quantified as portion of junction reads splicing from exon 16 of *Syngap1*. Reads with the *Syngap1* exon 16 splice donor site at the 3' end of exon 16 spliced to the 'VK' exon extension at the 5' end of exon 17 and non-VK splice acceptor 6bp downstream in all mice, whereas only the *Syngap1*$^{+/c.3583\text{-}9G>A}$ mice displayed a small fraction of cryptic splice acceptor site -7bp upstream. A two-way ANOVA revealed a significant effect of isoform, and interaction between isoform and genotype (Genotype $F (3, 9) = 3.494$; $p = 0.0631$, Isoform $F (2, 18) = 601.4$; $p < 0.0001$, Interaction $F (6, 18) = 2.741$; $p = 0.0452$, $n = 3\text{-}4$ samples/group) and Šídák's post hoc tests confirmed a significant increase in the cryptic splicing splice junction abundance in *Syngap1*$^{+/c.3583\text{-}9G>A}$ mice compared to wild-type littermates ($p = 0.0365$), but no significant decreases of other junctions ($p > 0.1212$).
(D) Exon 18 β/non-β alternative splicing ratio quantified as portion of junction reads splicing from exon 17 of *Syngap1*. Reads with the *Syngap1* exon 17 splice donor site at the 3' end of exon 17 spliced to the β exon extension at the 5' end of exon 18 and non-β splice acceptor 13bp downstream in all mice at similar ratios. A two-way ANOVA revealed a significant effect of isoform only (Genotype $F (3, 9) = 1.085$; $p = 0.4039$, Isoform $F (1, 9) = 670.7$; $p < 0.0001$, Interaction $F (3, 9) = 3.077$; $p = 0.0832$, $n = 3\text{-}4$ samples/group) and Šídák's post hoc tests confirmed no significant differences between groups.
(E) Exon 18 α1/α2/γ alternative splicing ratio quantified as portion of junction reads splicing to exon 20 of *Syngap1*. Reads with the *Syngap1* exon 20 splice acceptor site at the 5' end of exon 20 spliced from the α2 exon extension at the 3' end of exon 18, the α1 donor 1bp upstream, and from the γ donor at the 3' end of exon 19 in all mice at similar ratios. A two-way ANOVA revealed a significant effect of isoform only (Genotype $F (3, 9) = 0.1648$; $p = 0.9174$, Isoform $F (1.104, 9.936) = 287.2$; $p < 0.0001$, Interaction $F (6, 18) = 0.6083$; $p = 0.7206$, $n = 3\text{-}4$ samples/group) and Šídák's post hoc tests confirmed no significant differences between groups.
(F) N-terminal splicing ratio quantified as portion of junction reads splicing to exon 4 of *Syngap1*. Reads with the *Syngap1* exon 4 splice acceptor site at the 5' end of exon 4 spliced from exon 3 (A1/2/4 isoforms), exon 1 (A3), the B exon, the E exon, and the D exon in all mice, at varying levels. A two-way ANOVA revealed a significant effect of isoform, and interaction between isoform and genotype (Genotype $F (3, 9) = 1.364$; $p = 0.3419$, Isoform $F (1.929, 17.36) = 89.97$; $p < 0.0001$, Interaction $F (12, 36) = 2.651$; $p = 0.0118$, $n = 3\text{-}4$ samples/group) and Šídák's post hoc tests displayed trends of increased D isoform splicing in *Syngap1*$^{+/c.3583\text{-}9G>A}$ mice compared to wild-type littermates ($p = 0.0750$), but no significant decreases of other junctions ($p > 0.05$).
(G) N-terminal splice junction read abundance (RPMG) among junction reads splicing to exon 4 of *Syngap1*. A two-way ANOVA revealed a significant effect of isoform, genotype, and interaction between isoform and genotype (Genotype $F (3, 9) = 7.442$; $p = 0.0083$, Isoform $F (2.482, 22.33) = 102.0$; $p < 0.0001$, Interaction $F (12, 36) = 5.408$; $p < 0.0001$, $n = 3\text{-}4$ samples/group) and Šídák's post hoc tests confirmed significantly lower abundance of A1/2/4 isoform splicing reads in *Syngap1*$^{+/c.3583\text{-}9G>A}$ mice compared to wild-type littermates ($p = 0.0210$), as well as *Syngap1*$^{+/-}$ mice compared to wild-type littermates ($p = 0.0437$).

Supplementary Figure. 2

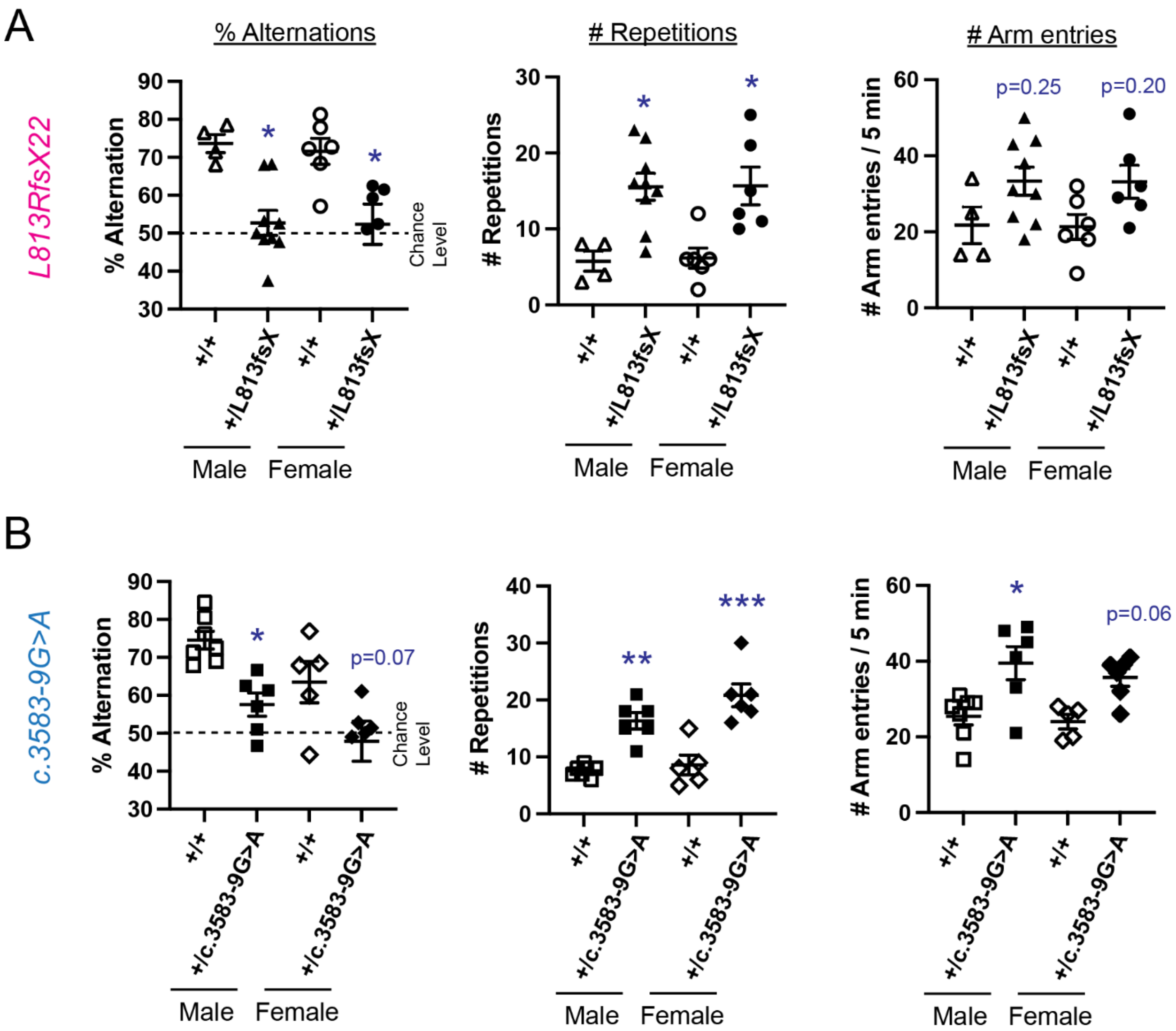


**Figure S2. Recapitulation of working memory deficits, repetitive behavior, and hyperactivity in SRID model mice (Data sorted by Sex)**

(A) Percent of spontaneous alternation (% alternation), the number of repetitive arm visits (# repetitions), and the number of arm visits (# arm entries) for wild-type (*Syngap1*$^{+/+}$) and *Syngap1*$^{+/L813RfsX22}$ are shown. One-way ANOVA followed by Tukey's post hoc test (Genotypes F (3,21) = 8.097; p = 0.0009 (% alternation), Genotypes F (3,21) = 8.085; p = 0.0009 (% repetitions), Genotypes F (3,21) = 2.720; p = 0.0703 (% alternation), n = 4-7 mice, *** p<0.001, ** p<0.01, * p< 0.05) was performed.

(B) % alternation, # repetitions, and # arm entries of for wild-type (*Syngap1*$^{+/+}$) and *Syngap1*$^{+/c.3583-9G>A}$ are shown. One-way ANOVA followed by Tukey's post hoc test (Genotype F (3, 20) = 8.325; p=0.0009 (% alternation), Genotype F (3, 20) = 20.02; p < 0.0001 (# repetitions), Genotype F (3, 20) = 6.685; p < 0.0001 (# arm entries), n = 4-7 mice, *** p<0.001, ** p<0.01, * < 0.05).

**Table S1. Top-regulated genes in whole-brain RNA-seq of *Syngap1*$^{+/c.3583-9G>A}$ mice and +/+ littermates.**

| ENSEMBL Gene ID | baseMean | log2FoldChange | lfcSE | stat | pvalue | padj | gene symbol |
|---|---|---|---|---|---|---|---|
| ENSMUSG00000067629 | 2923.401 | -0.6663 | 0.087803 | -7.00243 | 2.52E-12 | 2.96E-09 | Syngap1 |
| ENSMUSG00000025159 | 641.183 | 0.61346 | 0.099074 | 6.879794 | 5.99E-12 | 6.17E-09 | Mms19 |
| ENSMUSG00000078307 | 4494.512 | -0.33594 | 0.067046 | -5.0582 | 4.23E-07 | 3.45E-05 | AI593442 |
| ENSMUSG00000043439 | 1157.893 | -0.37525 | 0.081856 | -4.52488 | 6.04E-06 | 0.00032 | Epop |
| ENSMUSG00000026275 | 2047.27 | -0.36494 | 0.084028 | -4.26276 | 2.02E-05 | 0.000908 | Ppp1r7 |
| ENSMUSG00000009394 | 9787.411 | 0.28916 | 0.067259 | 4.235012 | 2.29E-05 | 0.001002 | Syn2 |
| ENSMUSG00000036062 | 4938.242 | -0.27172 | 0.064783 | -4.18109 | 2.90E-05 | 0.001226 | Phf24 |
| ENSMUSG00000023868 | 3889.187 | -0.30613 | 0.07592 | -3.93214 | 8.42E-05 | 0.003026 | Pde10a |
| ENSMUSG00000033065 | 8139.461 | 0.176768 | 0.04667 | 3.886674 | 0.000102 | 0.003522 | Pfkm |
| ENSMUSG00000039474 | 3132.889 | 0.397251 | 0.094393 | 3.827745 | 0.000129 | 0.004279 | Wfs1 |
| ENSMUSG00000024515 | 1114.135 | 0.336963 | 0.096692 | 3.766094 | 0.000166 | 0.00523 | Smad4 |
| ENSMUSG00000021614 | 364.0863 | 0.252596 | 0.105996 | 3.694525 | 0.00022 | 0.006661 | Vcan |
| ENSMUSG00000009406 | 1395.657 | 0.301485 | 0.077589 | 3.675656 | 0.000237 | 0.007089 | Elk1 |
| ENSMUSG00000015468 | 159.4931 | 0.328242 | 0.108255 | 3.670829 | 0.000242 | 0.007205 | Notch4 |
| ENSMUSG00000029223 | 11322.66 | 0.202907 | 0.054219 | 3.61991 | 0.000295 | 0.008565 | Uchl1 |
| ENSMUSG00000043671 | 1590.114 | -0.2286 | 0.10087 | -3.58266 | 0.00034 | 0.00963 | Dpy19l3 |
| ENSMUSG00000003949 | 4864.044 | -0.28106 | 0.075408 | -3.54803 | 0.000388 | 0.010822 | Hlf |
| ENSMUSG00000032579 | 162.6844 | 0.203972 | 0.107761 | 3.516166 | 0.000438 | 0.011896 | Hemk1 |
| ENSMUSG00000018405 | 296.5429 | -0.38206 | 0.106054 | -3.46081 | 0.000539 | 0.014139 | Mrm1 |
| ENSMUSG00000042423 | 573.163 | -0.24589 | 0.094185 | -3.3921 | 0.000694 | 0.017626 | Fbrs |
| ENSMUSG00000027894 | 12783.47 | 0.174154 | 0.054258 | 3.360899 | 0.000777 | 0.019442 | Slc6a17 |
| ENSMUSG00000051910 | 483.0625 | 0.245429 | 0.10066 | 3.288269 | 0.001008 | 0.024376 | Sox6 |
| ENSMUSG00000031833 | 3580.48 | -0.22958 | 0.075305 | -3.20455 | 0.001353 | 0.031113 | Mast3 |
| ENSMUSG00000039156 | 1337.171 | -0.26792 | 0.087652 | -3.13174 | 0.001738 | 0.038337 | Stim2 |
| ENSMUSG00000041890 | 738.6015 | -0.26525 | 0.097723 | -3.12617 | 0.001771 | 0.039044 | Git2 |
| ENSMUSG00000046447 | 39819.37 | -0.18787 | 0.060601 | -3.12551 | 0.001775 | 0.039053 | Camk2n1 |
| ENSMUSG00000019802 | 1851.024 | -0.26758 | 0.08482 | -3.11707 | 0.001827 | 0.039935 | Sec63 |
| ENSMUSG00000025889 | 3018.855 | 0.246697 | 0.078014 | 3.117923 | 0.001821 | 0.039935 | Snca |
| ENSMUSG00000042700 | 5287.667 | -0.24022 | 0.077389 | -3.1013 | 0.001927 | 0.041669 | Sipa1l1 |
| ENSMUSG00000017978 | 2832.53 | -0.23224 | 0.076012 | -3.07261 | 0.002122 | 0.045091 | Cadps2 |
| ENSMUSG00000001506 | 612.9428 | 0.346787 | 0.105052 | 3.051553 | 0.002277 | 0.047884 | Col1a1 |
| ENSMUSG00000036578 | 664.3525 | -0.22741 | 0.089835 | -3.02644 | 0.002475 | 0.051327 | Fxyd7 |
| ENSMUSG00000099083 | 669.7985 | 0.248561 | 0.099418 | 3.016066 | 0.002561 | 0.052652 | Atf7 |
| ENSMUSG00000060261 | 4876.012 | -0.20549 | 0.068361 | -2.98913 | 0.002798 | 0.056803 | Gtf2i |
| ENSMUSG00000025221 | 1528.742 | -0.2508 | 0.088106 | -2.98572 | 0.002829 | 0.057244 | Kcnip2 |
| ENSMUSG00000010803 | 11887.58 | -0.189 | 0.064855 | -2.98204 | 0.002863 | 0.057826 | Gabra1 |
| ENSMUSG00000027624 | 8794.183 | -0.16681 | 0.052746 | -2.98068 | 0.002876 | 0.058049 | Epb41l1 |
| ENSMUSG00000036353 | 695.7241 | 0.277751 | 0.089929 | 2.918946 | 0.003512 | 0.06874 | P2ry12 |
| ENSMUSG00000074923 | 964.0303 | -0.25608 | 0.089892 | -2.91596 | 0.003546 | 0.069113 | Pak6 |
| ENSMUSG00000075334 | 813.4046 | 0.300364 | 0.098406 | 2.900989 | 0.00372 | 0.072197 | Rprm |
| ENSMUSG00000089704 | 1316.547 | -0.26953 | 0.100193 | -2.88223 | 0.003949 | 0.075931 | Galnt2 |
| ENSMUSG00000115783 | 279.4018 | 0.327705 | 0.105822 | 2.851167 | 0.004356 | 0.082187 | AC161763.1 |
| ENSMUSG00000029608 | 6162.376 | -0.19047 | 0.066139 | -2.84454 | 0.004448 | 0.083578 | Rph3a |
| ENSMUSG00000031691 | 3287.516 | -0.18539 | 0.063357 | -2.83528 | 0.004579 | 0.085564 | Tnpo2 |
| ENSMUSG00000078161 | 339.6216 | -0.17381 | 0.102488 | -2.83522 | 0.004579 | 0.085564 | Erich3 |
| ENSMUSG00000044252 | 4987.782 | -0.16222 | 0.054565 | -2.8254 | 0.004722 | 0.087534 | Osbpl1a |
| ENSMUSG00000030729 | 8304.973 | 0.188803 | 0.06973 | 2.818134 | 0.00483 | 0.089042 | Pgm2l1 |
| ENSMUSG00000022351 | 1845.815 | 0.203072 | 0.075027 | 2.807399 | 0.004994 | 0.091565 | Sqle |
| ENSMUSG00000097156 | 551.1177 | -0.22326 | 0.094901 | -2.80508 | 0.00503 | 0.092049 | Gm3764 |
| ENSMUSG00000039637 | 921.7898 | -0.28914 | 0.09892 | -2.80345 | 0.005056 | 0.092217 | Coro7 |
| ENSMUSG00000022054 | 4215.059 | -0.19916 | 0.074511 | -2.79601 | 0.005174 | 0.093898 | Nefm |
| ENSMUSG00000032116 | 1195.852 | 0.232943 | 0.092582 | 2.793129 | 0.00522 | 0.094369 | Stt3a |
| ENSMUSG00000006005 | 940.1384 | -0.20807 | 0.101789 | -2.78846 | 0.005296 | 0.095329 | Tpr |
| ENSMUSG00000025795 | 1005.971 | -0.22659 | 0.084371 | -2.78175 | 0.005407 | 0.096742 | Rassf3 |

### Table S2. Top-regulated genes in whole-brain RNA-seq of *Syngap1*$^{+-}$ mice and +/+ littermates.

| ENSEMBL Gene ID | baseMean | log2FoldChange | lfcSE | stat | pvalue | padj | gene symbol |
|---|---|---|---|---|---|---|---|
| ENSMUSG00000041828 | 224.4626 | -0.55024 | 0.118633 | -4.38005 | 1.19E-05 | 0.000261 | Abca8a |
| ENSMUSG00000032796 | 107.0911 | -0.48849 | 0.119338 | -3.91515 | 9.03E-05 | 0.001264 | Lama1 |
| ENSMUSG00000053024 | 3285.634 | -0.29468 | 0.074284 | -3.82338 | 0.000132 | 0.001627 | Cntn2 |
| ENSMUSG00000071753 | 1071.017 | -0.37238 | 0.097406 | -3.80279 | 0.000143 | 0.001756 | C230004F18Rik |
| ENSMUSG00000039156 | 1337.171 | -0.30494 | 0.08137 | -3.76436 | 0.000167 | 0.002016 | Stim2 |
| ENSMUSG00000009112 | 1358.36 | -0.37364 | 0.097258 | -3.65659 | 0.000256 | 0.002981 | Bcl2l13 |
| ENSMUSG00000117098 | 516.9653 | -0.40686 | 0.110437 | -3.58405 | 0.000338 | 0.003803 | Gm49909 |
| ENSMUSG00000027883 | 254.4709 | -0.42435 | 0.119351 | -3.43511 | 0.000592 | 0.006333 | Gpsm2 |
| ENSMUSG00000053007 | 203.5276 | 0.410692 | 0.117616 | 3.421499 | 0.000623 | 0.006641 | Creb5 |
| ENSMUSG00000004637 | 565.0401 | 0.384862 | 0.106362 | 3.416118 | 0.000635 | 0.006767 | Wwox |
| ENSMUSG00000110051 | 252.7124 | -0.38401 | 0.114956 | -3.33503 | 0.000853 | 0.008861 | Gm45322 |
| ENSMUSG00000026074 | 2530.48 | -0.20424 | 0.062974 | -3.33141 | 0.000864 | 0.008972 | Map4k4 |
| ENSMUSG00000041133 | 1287.767 | -0.20546 | 0.064374 | -3.27335 | 0.001063 | 0.010954 | Smc1a |
| ENSMUSG00000094347 | 172.5456 | -0.39318 | 0.11927 | -3.24095 | 0.001191 | 0.012255 | Olfr784 |
| ENSMUSG00000015747 | 754.1997 | -0.31855 | 0.098614 | -3.22654 | 0.001253 | 0.012882 | Vps45 |
| ENSMUSG00000069171 | 1462.194 | 0.262492 | 0.078399 | 3.137062 | 0.001707 | 0.017512 | Nr2f1 |
| ENSMUSG00000019802 | 1851.024 | -0.23443 | 0.07642 | -3.1353 | 0.001717 | 0.017612 | Sec63 |
| ENSMUSG00000040785 | 15100.27 | -0.13511 | 0.045336 | -3.12525 | 0.001777 | 0.018209 | Ttc3 |
| ENSMUSG00000014353 | 816.4004 | 0.256879 | 0.080435 | 3.103956 | 0.00191 | 0.01956 | Tmem87b |
| ENSMUSG00000084319 | 1894.892 | 0.223008 | 0.07344 | 3.097035 | 0.001955 | 0.020016 | Tpt1-ps3 |
| ENSMUSG00000020328 | 286.9448 | 0.349082 | 0.116707 | 3.038053 | 0.002381 | 0.02436 | Nudcd2 |
| ENSMUSG00000090000 | 753.9249 | 0.280383 | 0.093656 | 3.037625 | 0.002385 | 0.024388 | Ier3ip1 |
| ENSMUSG00000044681 | 212.529 | -0.36232 | 0.118811 | -3.00197 | 0.002682 | 0.027392 | Cnpy1 |
| ENSMUSG00000026970 | 434.0037 | -0.36106 | 0.118798 | -2.99683 | 0.002728 | 0.027833 | Rbms1 |
| ENSMUSG00000058006 | 373.4865 | -0.34207 | 0.116569 | -2.98643 | 0.002823 | 0.028788 | Mdn1 |
| ENSMUSG00000021488 | 2368.883 | -0.21507 | 0.073882 | -2.96467 | 0.00303 | 0.030877 | Nsd1 |
| ENSMUSG00000038759 | 472.813 | -0.26187 | 0.090155 | -2.93777 | 0.003306 | 0.033636 | Nup205 |
| ENSMUSG00000031785 | 2423.376 | 0.18869 | 0.066589 | 2.930463 | 0.003385 | 0.034427 | Adgrg1 |
| ENSMUSG00000026034 | 659.2149 | -0.29295 | 0.100956 | -2.92966 | 0.003393 | 0.034505 | Clk1 |
| ENSMUSG00000024826 | 965.5626 | 0.313793 | 0.100745 | 2.91522 | 0.003554 | 0.036132 | Dpf2 |
| ENSMUSG00000038026 | 1050.452 | -0.22256 | 0.07725 | -2.91472 | 0.00356 | 0.036178 | Kcnj9 |
| ENSMUSG00000026384 | 3115.107 | 0.165618 | 0.059184 | 2.911391 | 0.003598 | 0.036555 | Ptpn4 |
| ENSMUSG00000041297 | 837.3305 | -0.28078 | 0.096806 | -2.89807 | 0.003755 | 0.038111 | Cdk13 |
| ENSMUSG00000021891 | 397.3236 | -0.31846 | 0.110383 | -2.89471 | 0.003795 | 0.038508 | Mettl6 |
| ENSMUSG00000045114 | 4119.293 | -0.1483 | 0.053937 | -2.89134 | 0.003836 | 0.038912 | Prrt2 |
| ENSMUSG00000055491 | 313.7231 | -0.32689 | 0.112859 | -2.88796 | 0.003877 | 0.039321 | Pprc1 |
| ENSMUSG00000029168 | 780.0819 | 0.263548 | 0.089446 | 2.886721 | 0.003893 | 0.039464 | Dpysl5 |
| ENSMUSG00000022075 | 2154.78 | 0.162714 | 0.057846 | 2.862658 | 0.004201 | 0.042563 | Rhobtb2 |
| ENSMUSG00000028245 | 416.2188 | 0.276846 | 0.098008 | 2.857576 | 0.004269 | 0.043225 | Nsmaf |
| ENSMUSG00000054452 | 9441.248 | 0.106797 | 0.036981 | 2.85582 | 0.004293 | 0.043451 | Aes |
| ENSMUSG00000078202 | 732.3926 | 0.268065 | 0.096156 | 2.83488 | 0.004584 | 0.04639 | Nrarp |
| ENSMUSG00000020263 | 912.549 | -0.22585 | 0.081868 | -2.83168 | 0.00463 | 0.046843 | Appl2 |
| ENSMUSG00000021772 | 1819.751 | 0.233273 | 0.08425 | 2.826963 | 0.004699 | 0.047524 | Nkiras1 |
| ENSMUSG00000032024 | 526.7112 | 0.281985 | 0.097251 | 2.80074 | 0.005099 | 0.051548 | Clmp |
| ENSMUSG00000003657 | 3092.239 | -0.18146 | 0.059399 | -2.79717 | 0.005155 | 0.052105 | Calb2 |
| ENSMUSG00000005338 | 11491.37 | -0.1297 | 0.048139 | -2.78011 | 0.005434 | 0.054906 | Cadm3 |
| ENSMUSG00000038128 | 5878.607 | -0.19947 | 0.07321 | -2.77753 | 0.005477 | 0.055327 | Camk4 |
| ENSMUSG00000048978 | 9750.312 | 0.137353 | 0.048445 | 2.77509 | 0.005519 | 0.05571 | Nrsn1 |
| ENSMUSG00000051910 | 483.0625 | -0.28673 | 0.104125 | -2.769 | 0.005623 | 0.056746 | Sox6 |
| ENSMUSG00000001524 | 230.4246 | 0.31717 | 0.114996 | 2.757556 | 0.005824 | 0.058735 | Gtf2h4 |
| ENSMUSG00000107495 | 227.149 | -0.32052 | 0.116096 | -2.75585 | 0.005854 | 0.059024 | Gm44215 |
| ENSMUSG00000030226 | 2474.986 | 0.183324 | 0.061369 | 2.746278 | 0.006028 | 0.060757 | Lmo3 |
| ENSMUSG00000030102 | 15665.86 | -0.22229 | 0.082268 | -2.74315 | 0.006085 | 0.061319 | Itpr1 |
| ENSMUSG00000009406 | 1395.657 | 0.200084 | 0.06697 | 2.729558 | 0.006342 | 0.063887 | Elk1 |
| ENSMUSG00000038070 | 108.0907 | -0.32056 | 0.119367 | -2.70985 | 0.006731 | 0.06779 | Cntln |
| ENSMUSG00000025582 | 7794.675 | -0.15556 | 0.059841 | -2.70063 | 0.006921 | 0.069677 | Nptx1 |
| ENSMUSG00000018405 | 296.5429 | -0.32668 | 0.115494 | -2.70017 | 0.00693 | 0.069751 | Mrm1 |
| ENSMUSG00000044452 | 539.2192 | -0.27017 | 0.09836 | -2.69722 | 0.006992 | 0.070351 | Zfp507 |
| ENSMUSG00000025576 | 4037.038 | -0.16688 | 0.064195 | -2.69668 | 0.007003 | 0.070444 | Rbfox3 |
| ENSMUSG00000025538 | 235.271 | -0.31164 | 0.114518 | -2.68935 | 0.007159 | 0.071989 | Sumf2 |
| ENSMUSG00000041774 | 256.6736 | 0.286016 | 0.107892 | 2.672059 | 0.007539 | 0.075783 | Ydjc |
| ENSMUSG00000020463 | 992.238 | -0.21553 | 0.07628 | -2.66598 | 0.007677 | 0.077145 | Ppp4r3b |
| ENSMUSG00000035206 | 592.573 | -0.26578 | 0.101293 | -2.65609 | 0.007905 | 0.079419 | Sppl2b |
| ENSMUSG00000105207 | 192.2732 | -0.32828 | 0.118604 | -2.65162 | 0.008011 | 0.080455 | Gm42927 |
| ENSMUSG00000026556 | 308.1257 | 0.313629 | 0.11859 | 2.632097 | 0.008486 | 0.085203 | Vangl2 |
| ENSMUSG00000062785 | 6215.709 | -0.15507 | 0.062769 | -2.62992 | 0.00854 | 0.085724 | Kcnc3 |
| ENSMUSG00000078897 | 111.4932 | 0.188068 | 0.091555 | 2.619387 | 0.008809 | 0.088392 | Gm4724 |
| ENSMUSG00000043557 | 724.6945 | -0.25973 | 0.098874 | -2.60738 | 0.009124 | 0.091497 | Mdga1 |
| ENSMUSG00000015647 | 139.5665 | -0.31215 | 0.110699 | -2.60585 | 0.009165 | 0.09188 | Lama5 |
| ENSMUSG00000061524 | 1373.619 | -0.24996 | 0.097713 | -2.60074 | 0.009302 | 0.093204 | Zic2 |
| ENSMUSG00000029416 | 521.4592 | 0.306272 | 0.108995 | 2.598183 | 0.009372 | 0.093873 | Slc15a4 |
| ENSMUSG00000049482 | 183.95 | 0.327766 | 0.119236 | 2.590571 | 0.009582 | 0.095917 | Ctu2 |
| ENSMUSG00000043770 | 208.0247 | 0.279192 | 0.109717 | 2.58875 | 0.009633 | 0.096397 | Gm12481 |
| ENSMUSG00000027167 | 318.0666 | 0.283136 | 0.112367 | 2.583391 | 0.009783 | 0.097878 | Elp4 |
| ENSMUSG00000058267 | 651.8682 | 0.214523 | 0.081913 | 2.57865 | 0.009919 | 0.099202 | Mrps14 |
| ENSMUSG00000027665 | 1789.591 | -0.26338 | 0.102463 | -2.57709 | 0.009963 | 0.09962 | Pik3ca |

### Table S3. Top-regulated genes in whole-brain RNA-seq of pooled *Syngap1* loss-of-function mice and +/+ littermates.

| ENSEMBL Gene ID | baseMean | log2FoldChange | lfcSE | stat | pvalue | padj | gene symbol |
|---|---|---|---|---|---|---|---|
| ENSMUSG00000039156 | 1337.171 | -0.24407 | 0.046464 | -5.59599 | 2.19E-08 | 2.81E-06 | Stim2 |
| ENSMUSG00000009406 | 1395.657 | 0.208963 | 0.04157 | 4.869753 | 1.12E-06 | 9.22E-05 | Elk1 |
| ENSMUSG00000054452 | 9441.248 | 0.121347 | 0.026779 | 4.553557 | 5.27E-06 | 0.00027 | Aes |
| ENSMUSG00000027883 | 254.4709 | -0.21246 | 0.04514 | -4.20479 | 2.61E-05 | 0.000535 | Gpsm2 |
| ENSMUSG00000018405 | 296.5429 | -0.16179 | 0.037332 | -3.90641 | 9.37E-05 | 0.001173 | Mrm1 |
| ENSMUSG00000004637 | 565.0401 | 0.180523 | 0.049485 | 3.846775 | 0.00012 | 0.001446 | Wwox |
| ENSMUSG00000043770 | 208.0247 | 0.167055 | 0.049577 | 3.826016 | 0.00013 | 0.001543 | Gm12481 |
| ENSMUSG00000038026 | 1050.452 | -0.16272 | 0.043844 | -3.80987 | 0.000139 | 0.001615 | Kcnj9 |
| ENSMUSG00000020263 | 912.549 | -0.16517 | 0.045275 | -3.73568 | 0.000187 | 0.002131 | Appl2 |
| ENSMUSG00000009112 | 1358.36 | -0.17048 | 0.049413 | -3.68724 | 0.000227 | 0.00256 | Bcl2l13 |
| ENSMUSG00000019802 | 1851.024 | -0.17471 | 0.047222 | -3.66791 | 0.000245 | 0.00276 | Sec63 |
| ENSMUSG00000053024 | 3285.634 | -0.16675 | 0.045795 | -3.66446 | 0.000248 | 0.002796 | Cntn2 |
| ENSMUSG00000038128 | 5878.607 | -0.16123 | 0.04541 | -3.63584 | 0.000277 | 0.003125 | Camk4 |
| ENSMUSG00000039637 | 921.7898 | -0.17076 | 0.049492 | -3.57323 | 0.000353 | 0.003975 | Coro7 |
| ENSMUSG00000028673 | 772.1169 | 0.166811 | 0.048074 | 3.533041 | 0.000411 | 0.00463 | Fuca1 |
| ENSMUSG00000044252 | 4987.782 | -0.10209 | 0.028953 | -3.52399 | 0.000425 | 0.004789 | Osbpl1a |
| ENSMUSG00000022391 | 5467.746 | 0.105991 | 0.030328 | 3.514741 | 0.00044 | 0.004958 | Rangap1 |
| ENSMUSG00000040785 | 15100.27 | -0.13291 | 0.039625 | -3.4743 | 0.000512 | 0.005765 | Ttc3 |
| ENSMUSG00000029053 | 4814.703 | -0.10242 | 0.030773 | -3.32478 | 0.000885 | 0.00995 | Prkcz |
| ENSMUSG00000036752 | 7065.12 | 0.103146 | 0.032193 | 3.311533 | 0.000928 | 0.01043 | Tubb4b |
| ENSMUSG00000107495 | 227.149 | -0.1454 | 0.048372 | -3.30107 | 0.000963 | 0.010823 | Gm44215 |
| ENSMUSG00000013629 | 472.1043 | 0.15397 | 0.046908 | 3.297927 | 0.000974 | 0.010937 | Cad |
| ENSMUSG00000026034 | 659.2149 | -0.13842 | 0.04942 | -3.29796 | 0.000974 | 0.010937 | Clk1 |
| ENSMUSG00000035245 | 302.9506 | 0.15432 | 0.049436 | 3.282157 | 0.00103 | 0.011564 | Eogt |
| ENSMUSG00000050608 | 1443.384 | 0.115615 | 0.037971 | 3.273701 | 0.001061 | 0.011911 | Minos1 |
| ENSMUSG00000029449 | 720.2359 | 0.142242 | 0.043973 | 3.265508 | 0.001093 | 0.012257 | Rhof |
| ENSMUSG00000044791 | 896.8124 | -0.13908 | 0.048896 | -3.26466 | 0.001096 | 0.01229 | Setd2 |
| ENSMUSG00000021488 | 2368.883 | -0.13369 | 0.044555 | -3.26153 | 0.001108 | 0.012418 | Nsd1 |
| ENSMUSG00000026970 | 434.0037 | -0.13646 | 0.045357 | -3.25718 | 0.001125 | 0.012606 | Rbms1 |
| ENSMUSG00000020160 | 235.5771 | -0.15264 | 0.049428 | -3.24735 | 0.001165 | 0.013045 | Meis1 |
| ENSMUSG00000029223 | 11322.66 | 0.115686 | 0.038049 | 3.238253 | 0.001203 | 0.013464 | Uchl1 |
| ENSMUSG00000020463 | 992.238 | -0.14187 | 0.043591 | -3.21411 | 0.001308 | 0.014644 | Ppp4r3b |
| ENSMUSG00000043557 | 724.6945 | -0.12399 | 0.049459 | -3.19242 | 0.001411 | 0.015779 | Mdga1 |
| ENSMUSG00000038759 | 472.813 | -0.1392 | 0.047927 | -3.18954 | 0.001425 | 0.015932 | Nup205 |
| ENSMUSG00000026839 | 552.2004 | 0.160791 | 0.048831 | 3.158296 | 0.001587 | 0.017725 | Upp2 |
| ENSMUSG00000028134 | 2252.802 | -0.13095 | 0.042479 | -3.13918 | 0.001694 | 0.018917 | Ptbp2 |
| ENSMUSG00000045994 | 2602.989 | -0.11122 | 0.03633 | -3.13503 | 0.001718 | 0.01918 | B3gat1 |
| ENSMUSG00000035206 | 592.573 | -0.12788 | 0.049332 | -3.12587 | 0.001773 | 0.019774 | Sppl2b |
| ENSMUSG00000003279 | 8293.032 | -0.11701 | 0.03909 | -3.11375 | 0.001847 | 0.020591 | Dlgap1 |
| ENSMUSG00000022756 | 1337.324 | 0.154907 | 0.047748 | 3.111633 | 0.001861 | 0.020732 | Slc7a4 |
| ENSMUSG00000050069 | 622.1054 | 0.143652 | 0.046143 | 3.109615 | 0.001873 | 0.020868 | Grem2 |
| ENSMUSG00000090000 | 753.9249 | 0.142682 | 0.049073 | 3.091942 | 0.001989 | 0.022136 | Ier3ip1 |
| ENSMUSG00000042523 | 2215.479 | -0.1156 | 0.038931 | -3.08939 | 0.002006 | 0.02232 | Dnal1 |
| ENSMUSG00000090136 | 352.2533 | 0.145544 | 0.04952 | 3.077128 | 0.00209 | 0.023243 | Gm10177 |
| ENSMUSG00000026596 | 1741.739 | -0.13324 | 0.048065 | -3.07461 | 0.002108 | 0.023432 | Abl2 |
| ENSMUSG00000030982 | 1185.124 | 0.139201 | 0.046108 | 3.073365 | 0.002117 | 0.023523 | 9030624J02Rik |
| ENSMUSG00000026074 | 2530.48 | -0.13163 | 0.047157 | -3.03622 | 0.002396 | 0.026598 | Map4k4 |
| ENSMUSG00000021451 | 1445.509 | -0.13264 | 0.047048 | -3.01565 | 0.002564 | 0.028441 | Sema4d |
| ENSMUSG00000024953 | 4212.177 | 0.10747 | 0.037373 | 3.002267 | 0.00268 | 0.029703 | Prdx5 |
| ENSMUSG00000029655 | 565.1538 | 0.124423 | 0.049316 | 3.000806 | 0.002693 | 0.029826 | N4bp2l2 |
| ENSMUSG00000037579 | 1594.892 | -0.12883 | 0.042622 | -2.99221 | 0.00277 | 0.030669 | Kcnh3 |
| ENSMUSG00000036568 | 1188.78 | -0.12418 | 0.047611 | -2.98227 | 0.002861 | 0.031672 | Bicral |
| ENSMUSG00000033161 | 19736.74 | -0.10819 | 0.036642 | -2.98103 | 0.002873 | 0.03179 | Atp1a1 |
| ENSMUSG00000048978 | 9750.312 | 0.107601 | 0.037648 | 2.97615 | 0.002919 | 0.032289 | Nrsn1 |
| ENSMUSG00000026319 | 1566.803 | -0.11746 | 0.039763 | -2.97471 | 0.002933 | 0.032431 | 2310035C23Rik |
| ENSMUSG00000062590 | 434.4995 | 0.150321 | 0.048815 | 2.967721 | 0.003 | 0.033166 | Armc9 |
| ENSMUSG00000097156 | 551.1177 | -0.12445 | 0.048887 | -2.96498 | 0.003027 | 0.033452 | Gm3764 |
| ENSMUSG00000014353 | 816.4004 | 0.143237 | 0.047469 | 2.921078 | 0.003488 | 0.038511 | Tmem87b |
| ENSMUSG00000044927 | 729.636 | 0.129376 | 0.044605 | 2.911316 | 0.003599 | 0.039722 | H1fx |
| ENSMUSG00000030516 | 2067.423 | -0.13358 | 0.046342 | -2.9062 | 0.003658 | 0.040364 | Tjp1 |
| ENSMUSG00000090223 | 4265.543 | -0.11976 | 0.040226 | -2.90571 | 0.003664 | 0.040414 | Pcp4 |
| ENSMUSG00000023868 | 3889.187 | -0.13147 | 0.045077 | -2.8851 | 0.003913 | 0.043114 | Pde10a |
| ENSMUSG00000020848 | 964.9516 | -0.12649 | 0.046098 | -2.88486 | 0.003916 | 0.043133 | Doc2b |
| ENSMUSG00000022748 | 158.9028 | 0.138469 | 0.048933 | 2.879697 | 0.003981 | 0.043831 | Cmss1 |
| ENSMUSG00000019907 | 1570.272 | -0.1112 | 0.041025 | -2.87762 | 0.004007 | 0.044107 | Ppp1r12a |
| ENSMUSG00000021244 | 1349.38 | -0.10509 | 0.038687 | -2.87448 | 0.004047 | 0.044517 | Ylpm1 |
| ENSMUSG00000022159 | 632.1918 | 0.137956 | 0.049401 | 2.847037 | 0.004413 | 0.048495 | Rab2b |
| ENSMUSG00000024847 | 1646.965 | 0.111181 | 0.041675 | 2.826487 | 0.004706 | 0.051651 | Aip |
| ENSMUSG00000071753 | 1071.017 | -0.17025 | 0.048702 | -2.82416 | 0.004741 | 0.052011 | C230004F18Rik |
| ENSMUSG00000039103 | 137.6118 | -0.10634 | 0.046565 | -2.82309 | 0.004756 | 0.052167 | Nexn |
| ENSMUSG00000079197 | 364.0376 | 0.122462 | 0.049346 | 2.79999 | 0.00511 | 0.056014 | Psme2 |
| ENSMUSG00000022723 | 227.8175 | -0.12095 | 0.041626 | -2.78803 | 0.005303 | 0.058087 | Crybg3 |
| ENSMUSG00000039048 | 376.7238 | -0.12944 | 0.049271 | -2.78426 | 0.005365 | 0.058746 | Foxred1 |
| ENSMUSG00000105230 | 185.8202 | 0.134928 | 0.049432 | 2.770595 | 0.005595 | 0.061209 | Gm42433 |
| ENSMUSG00000020923 | 1949.299 | -0.10834 | 0.041417 | -2.75848 | 0.005807 | 0.063504 | Ubtf |
| ENSMUSG00000027665 | 1789.591 | -0.14421 | 0.049511 | -2.75489 | 0.005871 | 0.064185 | Pik3ca |
| ENSMUSG00000020409 | 1112.591 | -0.12474 | 0.047322 | -2.74291 | 0.00609 | 0.06653 | Slu7 |

| | | | | | | | |
|---|---|---|---|---|---|---|---|
| ENSMUSG00000084319 | 1894.892 | 0.11591 | 0.045589 | 2.741455 | 0.006117 | 0.066804 | Tpt1-ps3 |
| ENSMUSG00000039474 | 3132.889 | 0.14909 | 0.049523 | 2.731797 | 0.006299 | 0.068749 | Wfs1 |
| ENSMUSG00000024998 | 406.3807 | -0.12079 | 0.049278 | -2.72902 | 0.006352 | 0.069309 | Plce1 |
| ENSMUSG00000041841 | 1661.581 | 0.102538 | 0.042746 | 2.728416 | 0.006364 | 0.069412 | Rpl37 |
| ENSMUSG00000039275 | 2170.658 | -0.10395 | 0.039631 | -2.72509 | 0.006428 | 0.070094 | Foxk2 |
| ENSMUSG00000057469 | 682.118 | -0.11753 | 0.043189 | -2.72436 | 0.006443 | 0.070225 | E2f6 |
| ENSMUSG00000074637 | 901.3402 | 0.117108 | 0.043446 | 2.723834 | 0.006453 | 0.070314 | Sox2 |
| ENSMUSG00000036893 | 541.2514 | -0.11923 | 0.049462 | -2.70193 | 0.006894 | 0.075069 | Ehmt1 |
| ENSMUSG00000006676 | 2130.417 | -0.12097 | 0.046008 | -2.6918 | 0.007107 | 0.077363 | Usp19 |
| ENSMUSG00000003037 | 814.2911 | -0.13765 | 0.048532 | -2.6909 | 0.007126 | 0.077547 | Rab8a |
| ENSMUSG00000078202 | 732.3926 | 0.129518 | 0.049212 | 2.684454 | 0.007265 | 0.079007 | Nrarp |
| ENSMUSG00000094347 | 172.5456 | -0.11579 | 0.043674 | -2.68006 | 0.007361 | 0.080025 | Olfr784 |
| ENSMUSG00000075028 | 399.4458 | 0.140746 | 0.049225 | 2.676758 | 0.007434 | 0.080766 | Prdm11 |
| ENSMUSG00000018076 | 1508.679 | -0.11333 | 0.047897 | -2.67476 | 0.007478 | 0.081203 | Med13l |
| ENSMUSG00000022377 | 2142.321 | -0.11177 | 0.042802 | -2.67473 | 0.007479 | 0.081203 | Asap1 |
| ENSMUSG00000061983 | 679.7515 | 0.130961 | 0.04832 | 2.672017 | 0.00754 | 0.08181 | Rps12 |
| ENSMUSG00000030302 | 13916.9 | -0.10227 | 0.039667 | -2.67118 | 0.007559 | 0.081988 | Atp2b2 |
| ENSMUSG00000021483 | 189.3851 | 0.116075 | 0.049377 | 2.670507 | 0.007574 | 0.082126 | Cdk20 |
| ENSMUSG00000074671 | 888.7839 | 0.110946 | 0.042529 | 2.663645 | 0.00773 | 0.083766 | Tspyl3 |
| ENSMUSG00000067629 | 2923.401 | -0.15874 | 0.049012 | -2.65912 | 0.007834 | 0.084843 | Syngap1 |
| ENSMUSG00000062866 | 748.103 | -0.11736 | 0.047508 | -2.65609 | 0.007905 | 0.085556 | Phactr2 |
| ENSMUSG00000058192 | 399.3893 | 0.131934 | 0.04935 | 2.651606 | 0.008011 | 0.086671 | Zfp846 |
| ENSMUSG00000018001 | 911.3443 | -0.11519 | 0.048462 | -2.64594 | 0.008146 | 0.08808 | Cyth3 |
| ENSMUSG00000021290 | 1412.645 | 0.105236 | 0.04307 | 2.632522 | 0.008475 | 0.091607 | 2010107E04Rik |
| ENSMUSG00000046364 | 1812.214 | 0.111458 | 0.048823 | 2.626272 | 0.008633 | 0.093246 | Rpl27a |
| ENSMUSG00000031691 | 3287.516 | -0.10987 | 0.043348 | -2.62579 | 0.008645 | 0.093349 | Tnpo2 |
| ENSMUSG00000008398 | 213.1505 | 0.11792 | 0.049515 | 2.623584 | 0.008701 | 0.093924 | Elk3 |
| ENSMUSG00000038705 | 445.7837 | -0.11552 | 0.047024 | -2.62324 | 0.00871 | 0.093988 | Gmeb2 |
| ENSMUSG00000004187 | 2989.606 | -0.11296 | 0.047751 | -2.62224 | 0.008735 | 0.094235 | Kifc2 |
| ENSMUSG00000004849 | 3891.48 | 0.114158 | 0.043551 | 2.617641 | 0.008854 | 0.095422 | Ap1s1 |
| ENSMUSG00000026384 | 3115.107 | 0.105348 | 0.039566 | 2.615521 | 0.008909 | 0.095986 | Ptpn4 |
| ENSMUSG00000039219 | 789.4158 | -0.10876 | 0.046729 | -2.61513 | 0.008919 | 0.096065 | Arid4b |
| ENSMUSG00000046707 | 972.9394 | -0.12309 | 0.049476 | -2.60848 | 0.009094 | 0.097919 | Csnk2a2 |
